# Characterisation of a novel ACE2-based therapeutic with enhanced rather than reduced activity against SARS-CoV2 variants

**DOI:** 10.1101/2021.03.17.435802

**Authors:** Mathieu Ferrari, Leila Mekkaoui, F. Tudor Ilca, Zulaikha Akbar, Reyisa Bughda, Katarina Lamb, Katarzyna Ward, Farhaan Parekh, Rajeev Karattil, Christopher Allen, Philip Wu, Vania Baldan, Giada Mattiuzzo, Emma M. Bentley, Yasuhiro Takeuchi, James Sillibourne, Preeta Datta, Alexander Kinna, Martin Pule, Shimobi C. Onuoha

## Abstract

The human angiotensin-converting enzyme 2 acts as the host cell receptor for SARS-CoV-2 and the other members of the *Coronaviridae* family SARS-CoV-1 and HCoV-NL63. Here we report the biophysical properties of the SARS-CoV-2 spike variants D614G, B.1.1.7 and B.1.351 with affinities to the ACE2 receptor and infectivity capacity, revealing weaknesses in the developed neutralising antibody approaches. Furthermore, we report a pre-clinical characterisation package for a soluble receptor decoy engineered to be catalytically inactive and immunologically inert, with broad neutralisation capacity, that represents an attractive therapeutic alternative in light of the mutational landscape of COVID-19. This construct efficiently neutralised four SARS-CoV-2 variants of concern. The decoy also displays antibody-like biophysical properties and manufacturability, strengthening its suitability as a first-line treatment option in prophylaxis or therapeutic regimens for COVID-19 and related viral infections.

## INTRODUCTION

The emergence of the severe acute respiratory syndrome coronavirus 2 (SARS-CoV-2) at the end of 2019^1^ has caused a major coronavirus disease (COVID-19) world-wide pandemic outbreak, totalling > 100 million confirmed cases and > 2 million associated deaths as of January 2021 (www.covid19.who.int). The rapid replication of SARS-CoV-2 has been shown in some patients to trigger an aggressive inflammatory response in the lung and acute respiratory disease syndrome (ARDS), leading to a cytokine release syndrome (CRS) due to the elevated expression of pro-inflammatory cytokines^2–4^. Similar to SARS-CoV-1^5^, this enveloped virus belongs to the β-coronavirus genus with a positive-strand RNA genome and utilises angiotensin-converting enzyme 2 (ACE2) as the receptor for host cell entry by binding to its glycoprotein Spike (S)^1,6^. The S protein is arranged as a trimeric complex of heterodimers composed of S1, containing the receptor binding domain (RBD) and S2, responsible for viral fusion and cell entry, which are generated from the proteolytical cleavage of the S precursor via furin in the host cell^6,7^.

Currently, more than 1100 monoclonal antibodies (mAb) against SARS-CoV-2 have been reported in the literature, with over 20 currently in clinical evaluation^8,9^. The antibodies LY-CoV555 and LY-CoV016 developed by Eli Lilly and Company, and the antibody cocktail REGN-COV2 (REGN10933 plus REGN10987) developed by Regeneron, were granted Emergency-Use Authorization (EUA) by the Food and Drug Administration (FDA). To maximise neutralisation capacity, most of the antibodies in development are directed towards the RBD, in order to disrupt interaction between the viral S-protein and ACE2^10^. These recombinant antibodies block viral entry by binding various epitopes on the RBD in a manner that fundamentally differs from the binding of the glycoprotein to ACE2 and are therefore susceptible to viral mutational escape.

Several variants have emerged carrying mutations in the S-protein, including in the RBD. Of note is the identification of the D614G (clade 20A) that has rapidly become the dominant strain globally^11^. Additional variants have also gained partial dominance in different regions of the globe. The variants A222V (clade 20A.EU1) and S477N (clade 20A.EU2) emerged in the summer of 2020 in Spain and have rapidly shown diffusion within Europe^12^. Recently, two new variants, clade 20B/501Y.V1, B.1.1.7 and clade 20C/501Y.V2, B.1.351, characterised by multiple mutations in S-protein have been associated with a rapid surge in COVID-19 cases in the UK and South Africa, respectively, and shown increased transmissibility and reduction of convalescent serum neutralisation capacity^13–15^. Finally, a variant that emerged in Brazil (B.1.1.28) contained mutational hallmarks of both the UK and South Africa variants, suggesting convergent evolution in SARS-CoV-2 due to similar selective pressures^16^. These variants have already been shown to impact on mAbs neutralisation potency^17,18^.

Receptor-based decoy strategies have successfully been employed in the clinic^19^-^21^, similarly, ACE2based decoy strategies have been proposed for COVID-19. A key advantage is that S-protein mutations which disrupt viral interaction with the ACE2 decoy, will by necessity decrease virulence thereby preventing meaningful escape by mutation. Previously described ACE2-based decoys include the soluble human catalytically active ACE2, repurposed from its initial development for treatment of non-COVID-19 ARDS^22^. Additionally, ACE2 mutants with enhanced affinity for the SARS-CoV-2 viral glycoprotein have also been described^23–25^. However, limitations of these approaches include short circulating half-life, activity over the renin/angiotensin system which may prevent its use in prophylaxis, and viral mutational escape which may be enabled by engineering of the S-protein targeting domain of ACE2.

With a view to eliminating the risk of mutational escape, eliminating the physiological effects on the renin/angiotensin system and increasing circulating half-life, we generated a catalytically inactive ACE2 receptor decoy fused to a human Fc domain further engineered to bear minimal immuno-modulatory activity. This molecule has shown complete lack of enzymatic activity and natural substrate sequestration, with no residual engagement to human FcγRs, adopting a set of Fc mutations reported to preserve long half-life and FcRn interaction^26^. The construct showed broad neutralising capacity with proven activity towards ACE2-tropic viruses, including the SARS-CoV-2 variants of concern B.1.1.7 and B.1.351, with improved consistency and resistance to viral mutational escape compared to leading monoclonal antibody therapeutics. Additionally, we report the biophysical characterisation and ACE2 affinity measurements for the D614G, B.1.1.7 and B.1.351 SARS-CoV-2 S1 variants, with links to infective potency in a pseudotyped vector setting, with direct comparison to approved COVID-19 monoclonal antibodies.

## RESULTS

### Biophysical characterisation of SARS-CoV-2 spike variants

We first explored the binding kinetic between SARS-CoV-2 S1 and ACE2. Inhouse purified recombinant S1 domains from WT, D614G, B.1.1.7 and B. 1.351 variants demonstrated similar properties to commercially sourced S1 WT protein (**Supplementary Figure 1A and 1B**). Interestingly, the WT and D614G variants displayed a similar thermal unfolding profile with the first transition event (Tm) at 42.9 and 42.2°C, respectively, while the B.1.1.7 and B.1.351 resulted in a 6.9 and 11.5°C increase compared to S1 WT, respectively (**Supplementary Figure 1C**).

The binding affinity of the spike variants for the ACE2 receptor was assessed by surface plasmon resonance (SPR) using the recombinant S1 domains to allow for a monovalent binding interaction. The SARS-CoV-2 S1 WT, D614G and B.1.351 displayed overall similar kinetic affinities, although the latter showed a 1.5-fold slower off-rate (*k_d_*) compared to WT S1, which was compensated by a slightly slower on-rate (*k_a_*). The B.1.1.7 S1 variant however, showed a > 3-fold increase in affinity compared to WT, mainly driven by a 3.8-fold slower *k_d_* (**Figure 1A** and **Table 1**).

**Figure 1.**
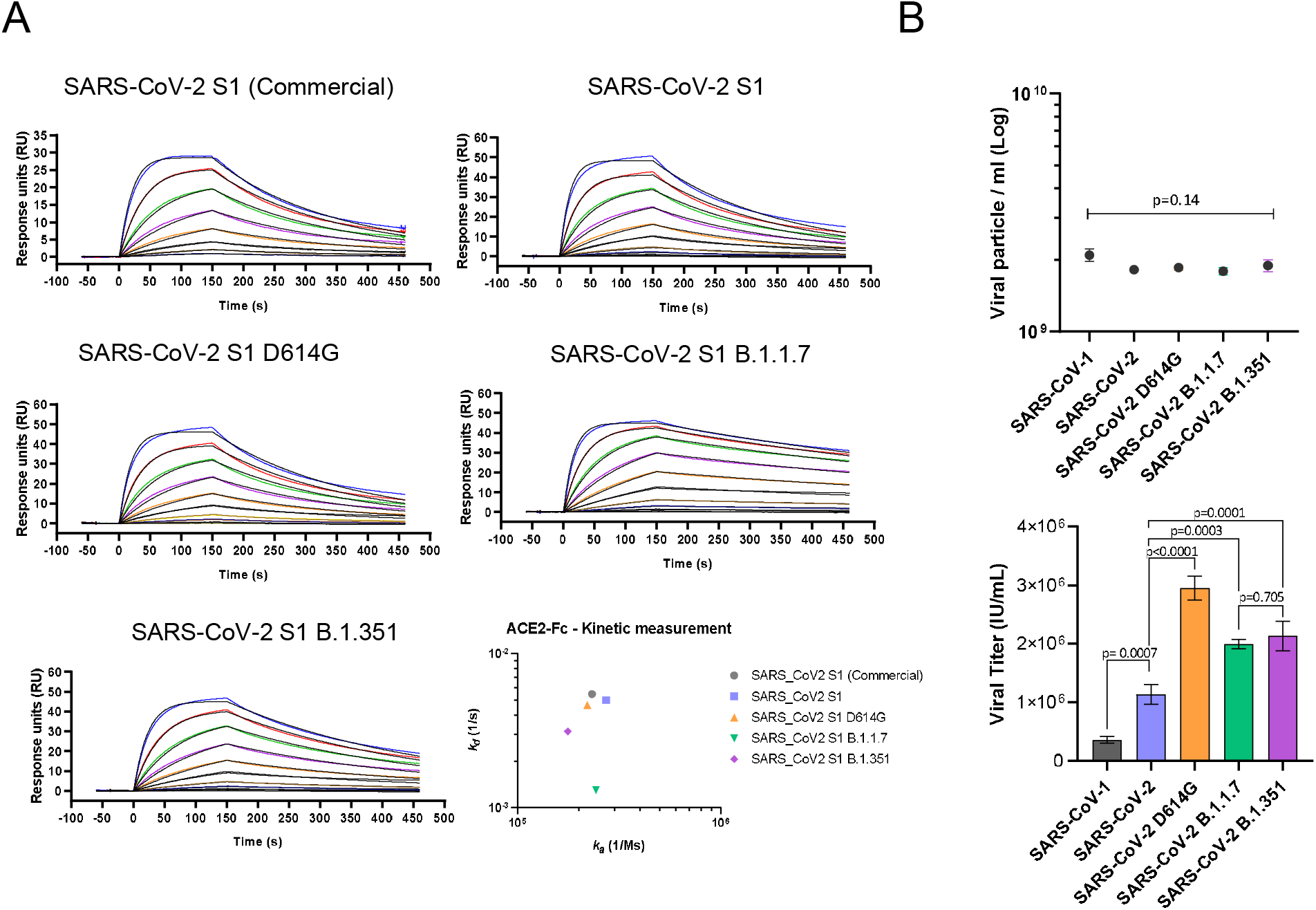
Biophysical characterisation of SARS-CoV-2 S1 variants. A) Kinetic affinity of catalytically active ACE2-Fc with SARS-CoV-2 S1 WT, D614G, B1.1.7 and B.1.351. All sensograms fitted with 1:1 Langmuir binding model. Analyte starting concentration 250 nM with 2-fold serial dilutions. 3-fold higher affinity detected for S1 of B.1.1.7 variant. B) Infectious viral titre (left) and p24 ELISA determined physical particle number (right) comparison of SARS-CoV-1, SARS-CoV-2, SARS-CoV-2 D614G, SARS-CoV-2 B1.1.7 and SARS-CoV-2 B1.351 pseutotyped vectors, showing comparable particle concentration but diverse infectivity capacity (Mean ± SD). One way ANOVA Dunnett’s multiple comparison (F=105.2, df=4, 10).

**Table 1.** Kinetic affinities.

| Clone | Spike S1 domain | 1:1 binding $k_a$<br>(1/Ms) | $k_d$ (1/s) | $K_D$ (M) | Two-state<br>$K_D$ (M) | Affinity fold change<br>(relative to<br>SARS-CoV-2 $K_D$) |
| --- | --- | --- | --- | --- | --- | --- |
| ACE2-Fc<br>active | SARS-CoV-1 | 4.17E+05 | 1.44E-02 | 3.45E-08 | 1.98E-08 |  |
|  | SARS-CoV-2 | 2.73E+05 | 5.00E-03 | 1.83E-08 |  | 1 |
|  | SARS-CoV-2 D614G | 2.20E+05 | 4.64E-03 | 2.11E-08 |  | -1.15 |
|  | SARS-CoV-2 B.1.1.7 | 2.43E+05 | 1.30E-03 | 5.33E-09 |  | 3.43 |
|  | SARS-CoV-2 B.1.351 | 1.77E+05 | 3.13E-03 | 1.78E-08 |  | 1.03 |
|  | HCoV-NL63 | 2.97E+03 | 3.55E-03 | 1.20E-06 |  |  |
| ACE2(HH:NN)<br>Fc | SARS-CoV-1 | 4.93E+05 | 1.31E-02 | 2.65E-08 | 9.31E-09 |  |
|  | SARS-CoV-2 | 2.72E+05 | 3.48E-03 | 1.28E-08 |  | 1 |
|  | SARS-CoV-2 D614G | 2.28E+05 | 3.18E-03 | 1.39E-08 |  | -1.09 |
|  | SARS-CoV-2 B.1.1.7 | 2.91E+05 | 1.01E-03 | 3.48E-09 |  | 3.68 |
|  | SARS-CoV-2 B.1.351 | 1.76E+05 | 2.39E-03 | 1.36E-08 |  | -1.06 |
|  | HCoV-NL63 | 4.00E+03 | 3.34E-03 | 8.35E-07 |  |  |
| ACE2(HH:NN)<br>Fc LALA-PG | SARS-CoV-1 | 5.05E+05 | 1.39E-02 | 2.75E-08 | 9.69E-09 |  |
|  | SARS-CoV-2 | 2.85E+05 | 3.27E-03 | 1.15E-08 |  | 1 |
|  | SARS-CoV-2 D614G | 2.40E+05 | 2.99E-03 | 1.24E-08 |  | -1.08 |
|  | SARS-CoV-2 B.1.1.7 | 2.83E+05 | 9.29E-04 | 3.28E-09 |  | 3.51 |
|  | SARS-CoV-2 B.1.351 | 1.72E+05 | 2.21E-03 | 1.28E-08 |  | -1.11 |
|  | HCoV-NL63 | 3.72E+03 | 3.49E-03 | 9.38E-07 |  |  |
| LY-CoV555 | SARS-CoV-1 | N/A | N/A | N/A |  |  |
|  | SARS-CoV-2 | 2.64E+05 | 9.96E-04 | 3.77E-09 |  | 1 |
|  | SARS-CoV-2 D614G | 2.11E+05 | 8.80E-04 | 4.17E-09 |  | -1.11 |
|  | SARS-CoV-2 B.1.1.7 | 2.22E+05 | 9.08E-04 | 4.08E-09 |  | -1.08 |
|  | SARS-CoV-2 B.1.351 | 1.68E+05 | 7.60E-03 | 4.51E-08 |  | -11.96 |
|  | HCoV-NL63 | N/A | N/A | N/A |  |  |
| REGN10933 | SARS-CoV-1 | N/A | N/A | N/A |  |  |
|  | SARS-CoV-2 | 1.00E+06 | 1.54E-03 | 1.54E-09 |  | 1 |
|  | SARS-CoV-2 D614G | 8.27E+05 | 1.25E-03 | 1.51E-09 |  | 1.02 |
|  | SARS-CoV-2 B.1.1.7 | 8.81E+05 | 1.40E-03 | 1.59E-09 |  | -1.03 |
|  | SARS-CoV-2 B.1.351 | 2.47E+05 | 8.64E-03 | 3.49E-08 |  | -22.67 |
|  | HCoV-NL63 | N/A | N/A | N/A |  |  |
| REGN10987 | SARS-CoV-1 | N/A | N/A | N/A |  |  |
|  | SARS-CoV-2 | 1.16E+06 | 9.36E-03 | 8.07E-09 |  | 1 |
|  | SARS-CoV-2 D614G | 8.88E+05 | 8.12E-03 | 9.14E-09 |  | -1.13 |
|  | SARS-CoV-2 B.1.1.7 | 5.39E+05 | 4.51E-03 | 8.36E-09 |  | -1.04 |
|  | SARS-CoV-2 B.1.351 | 3.64E+05 | 4.25E-03 | 1.17E-08 |  | -1.45 |
|  | HCoV-NL63 | N/A | N/A | N/A |  |  |

To assess the infectivity conferred by the SARS-CoV-2 spike variants, we engineered replication deficient lentiviral vectors pseudotyped with the WT glycoprotein or carrying the D614G, B.1.1.7 and B.1.351 mutations, alongside SARS-CoV-1. Although all pseudotyped vectors showed equivalent physical particle concentrations, as measured by p24 ELISA, they exhibited vastly different infectivity capacity (**Figure 1C**). SARS-CoV-1 resulted in the lowest viral titre with a reduction of 3.2-fold in infectious units (IU)/ml compared to WT SARS-CoV-2. The SARS-CoV-2 D614G variant was instead the most efficient with 2.6-fold higher viral titre compared to WT. B.1.1.7 and B.1.351 showed 1.8 and 1.9-fold higher viral titres, compared to WT SARS-CoV-2, respectively (**Figure 1C**).

### Catalytically inactive ACE2 -Fc fusion with streamlined purification

The extracellular domain of human ACE2 (aa 18-740, Uniprot Q9BYF1) was fused to the human IgG1 Fc via the human IgG1 hinge region to allow for homodimer stabilisation (**Figure 2A**). The ACE2 domain used included both the zinc metallopeptidase and the collectrin domains to allow full receptor representation. The Fc domain was included to improve circulating half-life and to capitalise on the streamlined antibody purification processes. In order to generate an inert receptor decoy, the catalytic site of the enzyme was mutated at residues 374 (H374N) and 378 (H378N), termed HH:NN, to inhibit enzymatic activity and prevent conversion of the Angiotensin 1-8 (Ang II) substrate to Angiotensin 1-7. This mutation is predicted to remove interaction with zinc ions (Zn^+2^) mediated by the two original Histidine (His) residues, with a spatially conservative mutation (**Figure 2B**).

**Figure 2.**
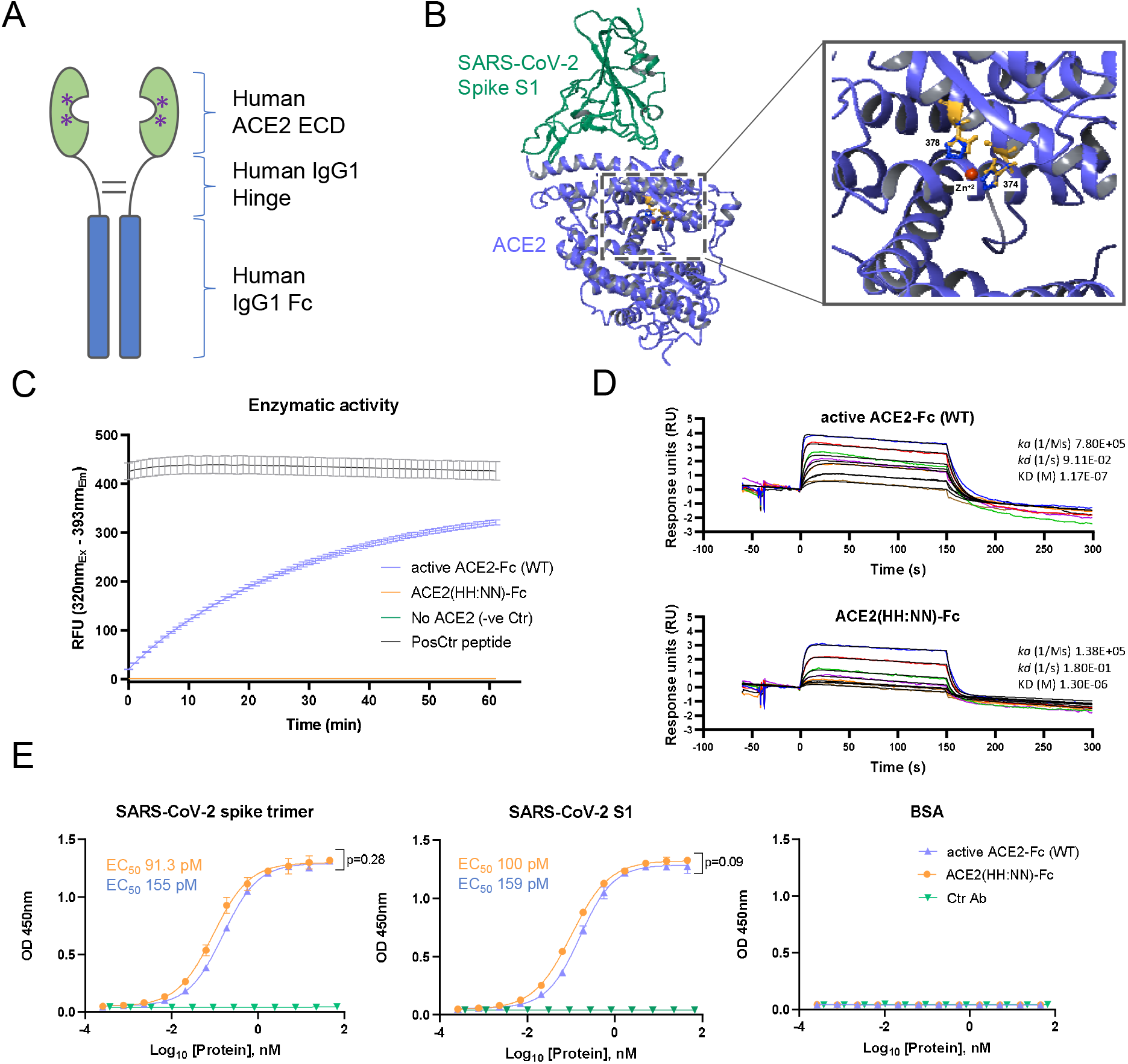
Characterisation of ACE2-Fc receptor decoy. A) Schematic representation of ACE2-Fc molecule with a streamlined antibody-like expression/purification process and biophysical characterisation. B) 3D structure of SARS-CoV-2 spike S1 domain (green) in complex with ACE2 (blue) (PDB 6M0J). Inset, zoomed section of the ACE2 catalytic site showing the H374 and H378 residues (blue) in complex with Zn (red), and the H374N and H378N mutations (orange). C) Enzymatic activity of active (blue) and H374N, H378N mutated (orange) ACE2-Fc using Mca-APK(Dnp) fluorogenic peptide (Mean ± SD). D) Binding kinetics of active (top) and inactive (bottom) ACE2-Fc with Ang II. E) ELISA of SARS-CoV-2 active spike trimer (left) or S1 domain (centre) against WT active and ACE2(HH:NN)-Fc, showing comparable binding capacity. No binding detected with control antigen (right) or negative control antibody (Mean ± SD). EC_50_ = half maximal effective concentration. Unpaired t-test of AUC (left t=1.086, df=24; centre t=1.79, df=24).

We first set out to confirm inactivation of the ACE2 component. *In vitro* testing using a fluorogenic substrate for ACE2, Mca-APK(Dnp), showed complete abrogation of enzymatic activity for the ACE2-Fc construct carrying the HH:NN mutation, while the wild-type (WT) active ACE2-Fc molecule was able to efficiently process the peptide (**Figure 2C**). Furthermore, the kinetic interaction of both WT and mutated ACE2 domains for their natural substrate Ang II was investigated using SPR. Both constructs interacted with the substrate, however, the mutated ACE2 was characterised by a slower on-rate (*k_a_* 7.80E+05 vs.1.38E+05, for active and HH:NN ACE2, respectively) and a faster off-rate *(k_d_* 9.11E-02 vs. 1.80E-01, for active and HH:NN ACE2, respectively), culminating in a final affinity (K_D_) of 1.3 μM for the ACE2 HH:NN vs. 117 nM for the WT active ACE2 (**Figure 2D**).

We next explored whether the ACE2 mutations impacted SP binding. Both WT and mutated ACE2 showed comparable binding capacity for recombinant SARS-CoV-2 full S trimer and S1 domain by ELISA (**Figure 2E**). SPR measurements of kinetic interaction for the S1 domain of SARS-CoV-2 showed comparable kinetic profiles between active WT and HH:NN ACE2 (**Table 1**), further suggesting the preservation of an unaltered Spike binding domain.

### Engineered Fc domain with abrogated FcγR engagement

To overcome the risk of activating the host immune system, thus exacerbating the hyperinflammatory response often associated with severe COVID-19 development^27^, the Fc domain was engineered to remove FcγR interactions. The well-established L234A/L235A (LALA)^28^ mutations of the CH2 domain and the LALA combination with P329G (LALA-PG)^29^ were introduced in the human IgG1 Fc portion of the ACE2-Fc fusion protein.

We first investigated the expression yields of the ACE2(HH:NN) with WT Fc, LALA Fc and LALA-PG Fc and ACE2 domain activity. All constructs showed comparable expression and purification efficiencies using protein A affinity chromatography (**Supplementary Figure 2**). Mutations on the Fc domain did not affect the binding capacity of ACE2 for SARS-CoV-2 S-protein and all three versions showed highly comparable dose/response curves to recombinant SARS-CoV-2 full S trimer or S1 domain by ELISA (**Figure 3A**). Similarly, all three variants were able to bind SupT1 cell lines expressing SARS-CoV-2 full S trimer as a transmembrane protein (**Figure 3B**), further confirming binding capacity for the glycoprotein in a more physiological environment.

**Figure 3.**
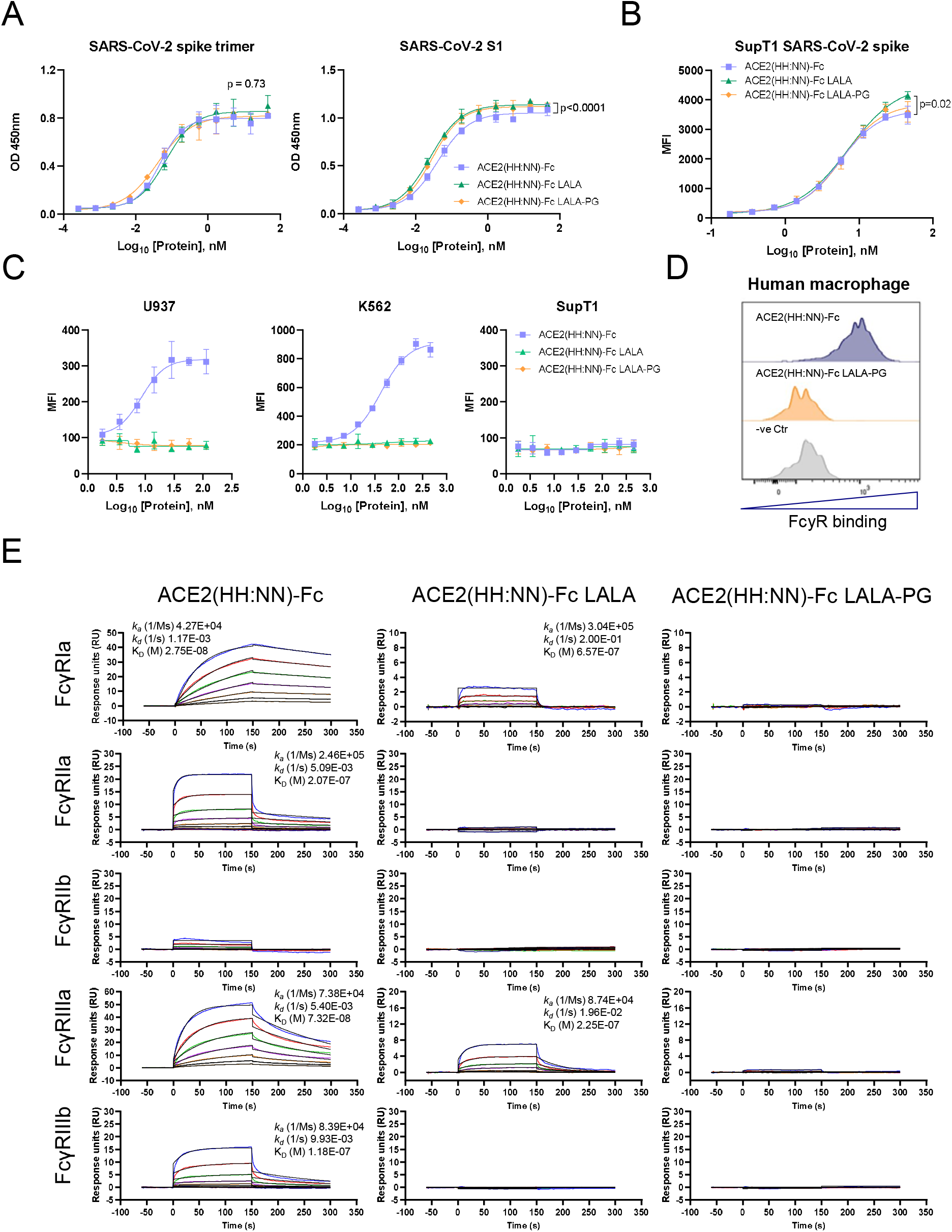
Characterisation of Fc effector functions. A) ELISA of SARS-CoV-2 active spike trimer (left) or S1 domain (right) with ACE2(HH:NN) WT Fc (blue), LALA Fc (green) or LALA-PG Fc (orange), showing comparable binding capacity (Mean ± SD). One way ANOVA of AUC with Turkey’s multiple comparison (left F=0.3121, df=2, 72; right F=34.17, df=2, 72, compared to blue). B) Binding capacity on SupT1 cell line expressing SARS-CoV-2 full length spike, by flow cytometry with ACE2(HH:NN) WT Fc (blue), LALA Fc (green) or LALA-PG Fc (orange). (Mean ± SD). One way ANOVA of AUC with Turkey’s multiple comparison (blue vs green, F=3.986, df=2, 54). C) Fc-mediated binding capacity to U937, K562 and SupT1 of ACE2(HH:NN) WT Fc (blue), LALA Fc (green) or LALA-PG Fc (orange), detected with biotinylated SARS-CoV-2 S1 and streptavidin conjugated secondary agent. No binding was detected with ACE2-Fc constructs carrying the LALA or LALA-PG mutations (Mean ± SD). D) Representative flow cytometry of Fc-mediated binding of ACE2(HH:NN) WT Fc (blue) and LALA-PG Fc (orange) on human monocyte-derived M1 macrophages. No binding detected with Fc carrying the LALA-PG mutation (n=4). E) SPR binding kinetic of ACE2(HH:NN) WT Fc, LALA Fc or LALA-PG Fc on human FcγRIa, FcγRIIa, FcγRIIb, FcγRIIIa and FcγRIIIb. LALA-PG mutations mediated a complete abrogation of FcγR interaction. Sensograms fitted with 1:1 Langmuir binding model.

Next, we investigated the residual interaction of the engineered Fc domains for human FcγRI, FcγRII and FcγRIII on K562, U937 and SupT1 human cell lines. K562 are reported to express RNA for FcγRIIa and IIIa/b, while U937 express FcγRIa/b, IIa/b and IIIb (source www.proteinatlas.org, v20.0). A reverse flow cytometry detection assay, using biotinylated SARS-CoV-2 S1 as secondary reagent, demonstrated that the ACE2 construct with WT Fc efficiently bound both K562 and U937 in a dose-dependent manner (**Figure 3C**). No binding was detected with either LALA or LALA-PG Fc mutations. SupT1 cells are not reported to express FcγR on the membrane and, consequently, failed to show binding events with the tested molecules. Equally, human M1 polarised monocyte-derived macrophages (MDM) from healthy donors, showed strong interaction with the ACE2 carrying WT Fc, while no detectable engagement was obtained with the LALA-PG Fc mutation (**Figure 3D**).

Binding affinities of WT, LALA and LALA-PG ACE2(HH:NN)-Fc variants for human FcγRs were tested via SPR. ACE2(HH:NN)-Fc showed strong interaction with FcγRIa and IIIa (27.5 nM and 73.2 nM, respectively) and reduced binding affinity for FcγRIIa and IIIb (207 nM and 118 nM, respectively). The LALA mutation still maintained residual binding to the FcγRIa and IIIa (657 nM and 225 nM, respectively) but no detectable binding to the remainder of the receptors. The LALA-PG mutation, however, showed a complete abrogation of FcγR binding, suggesting a more silent immunomodulatory profile (**Figure 3E**).

### Analysis of ACE2 decoy cross-reactivity and spike binding affinity

Binding specificity and cross-reactivity of the ACE2(HH:NN)-Fc LALA-PG construct was assessed using a cell-based protein microarray assay, screening 5477 full length plasma membrane and cell surface-tethered human secreted proteins, 371 human heterodimers, and the SARS-CoV-2 S-protein (**Supplementary Table 1**). The test construct showed strong specific binding to the target protein SARS-CoV-2 S, while no other interaction was detected across the comprehensive panel of human protein (**Figure 4A**). An Fc LALA-PG only construct with the ACE2 domain omitted did not display any interaction with SARS-CoV-2 S-protein or any other target tested. The control fusion protein CTLA4-hFc instead, showed strong interaction for its predicted target CD86, and the FcγRIa, due to the presence of a WT IgG1 Fc domain. A secondary anti-human Fc antibody interaction with human IgG3 was detected across all conditions tested (**Figure 4A**).

**Figure 4.**
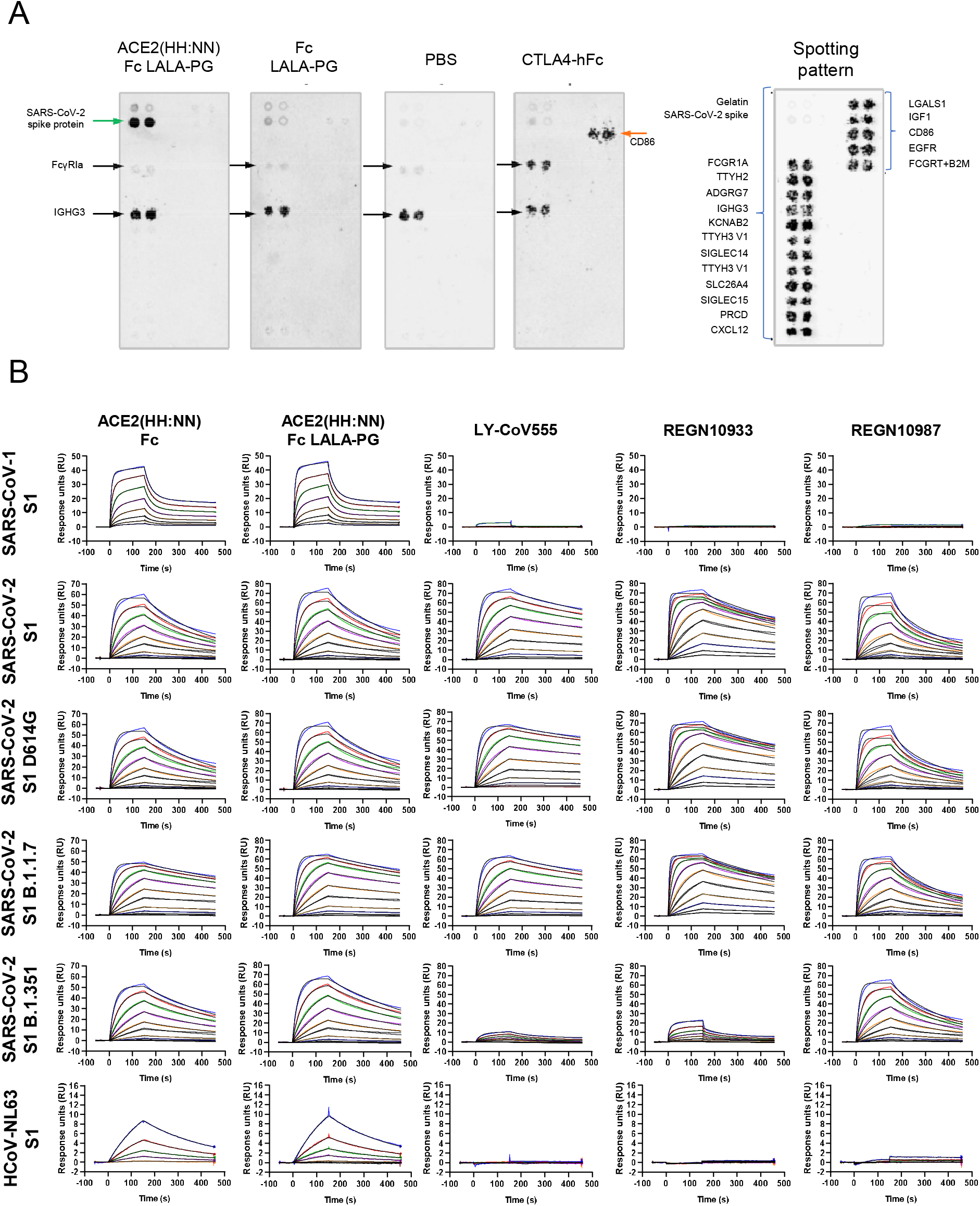
ACE2(HH:NN)-Fc LALA-PG specificity. A) Cell microarray screening of human cell-membrane proteome with ACE2(HH:NN)-Fc LALA-PG, control Fc (LALA-PG), CTLA4-hFc or PBS. Depicted is a selection of antigens (key legend on the right panel). ACE2(HH:NN)-Fc LALA-PG shows strong specific interaction with SARS-CoV-2 spike protein only. B) SPR binding kinetics of ACE2(HH:NN) WT Fc, LALA-PG Fc, LY-CoV555, REGN10933 and REGN10987 against SARS-CoV-1, SARS-CoV-2 variants (WT, D614G, B1.1.7 and B.1.351) and HCoV-NL63 S1 domains. ACE2(HH:NN) Fc and ACE2(HH:NN) Fc LALA-PG were able to efficiently bind all spike protein tested. All sensograms were fitted with Langmuir 1:1 binding model, except for SARS-CoV-1 S1 kinetics which were fitted with two-state kinetics. 2-fold serial dilutions starting from 250 nM (500 nM for HCoV-NL63 S1).

As this receptor decoy has the potential to bind S glycoproteins of viruses that utilise ACE2 as host-cell receptor, binding kinetics were generated for the S1 spike domain of SARS-CoV-1, SARS-CoV-2, SARS-CoV-2 D614G, B.1.1.7 and B.1.351 variants, and HCoV-NL63, comparing to the leading anti-SARS-CoV-2 antibodies LY-CoV555^30^, REGN10933 and REGN10987^31^. The ACE2-Fc fusion constructs mediated specific interaction towards all spike proteins tested, while the monoclonal antibodies showed specificity only for the SARS-CoV-2 related S-proteins (**Figure 4B**). The ACE2(HH:NN)-Fc and ACE2(HH:NN)-Fc LALA-PG showed comparable affinities for the tested S1 domains, confirming no effect of the Fc mutations on ACE2 binding (**Figure 4B and Table 1**). Similar to the active ACE2-Fc, the inactive ACE2-Fc constructs also displayed a 3-fold enhanced affinity for the SARS-CoV-2 S1 B.1.1.7. While the monoclonal antibody REGN10987 maintained a similar affinity for the SARS-CoV-2 S1 variants tested, the LY-CoV555 and REGN10933 were dramatically affected by the B.1.351 variant with 12- and 23-fold reduction in affinity compared to S1 WT, respectively (**Figure 4B** and **Table 1**).

### In vitro neutralisation of SARS-CoV-2 variants of concern

We first assessed the neutralisation capacity of our decoy receptor against live replicating SARS-CoV-2. Both ACE2(HH:NN)-Fc and ACE2(HH:NN)-Fc LALA-PG showed comparable neutralisation efficiency for live SARS-CoV-2 virus *in vitro,* with half maximal neutralisation titres (NT_50_) values of 5.2 and 4.1 nM, respectively, providing evidence of potent therapeutic activity (**Figure 5A**).

**Figure 5.**
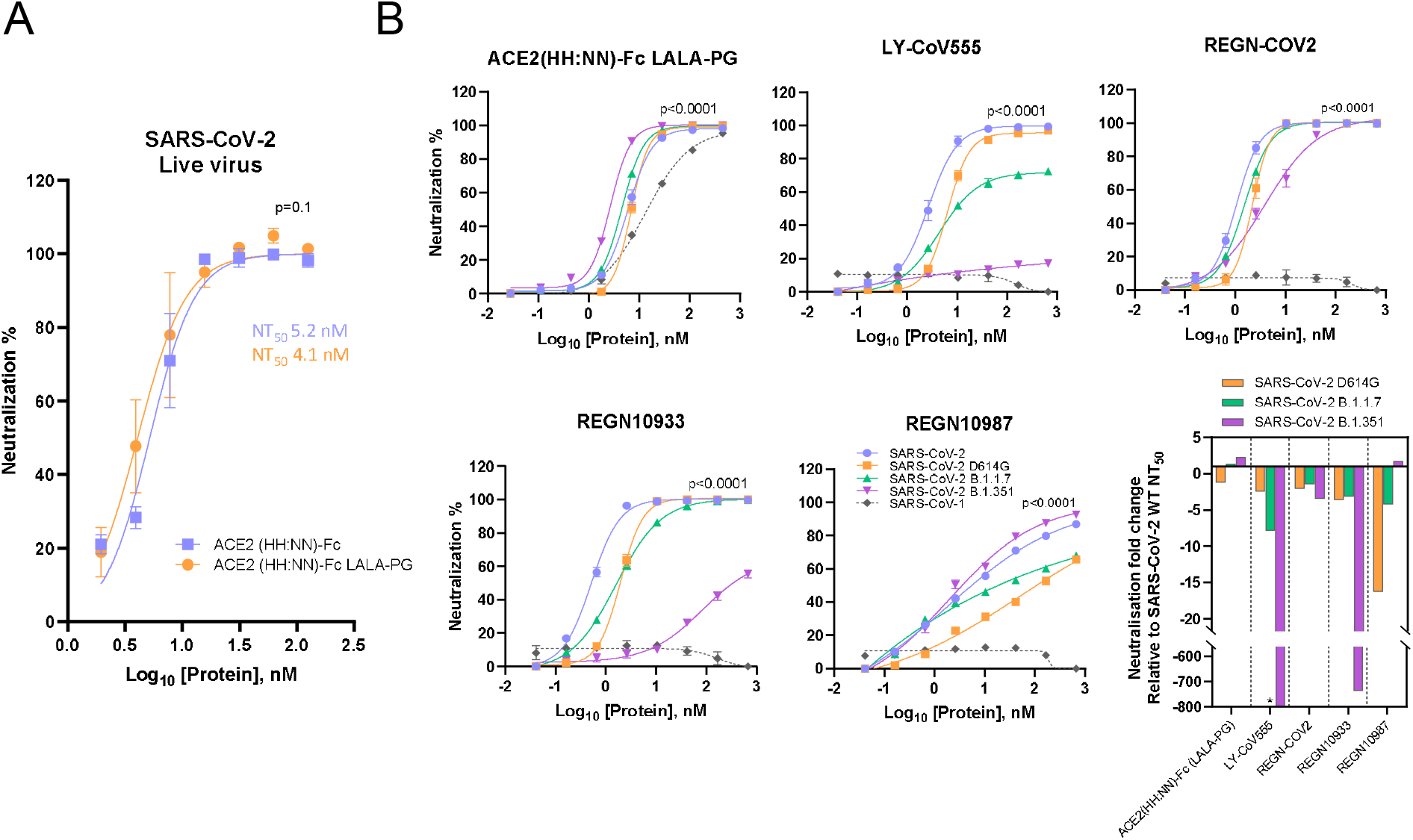
SARS-CoV-2 neutralisation efficiency. A) Neutralisation assay of live SARS-CoV-2 virus with ACE2(HH:NN) WT Fc (blue) and LALA-PG Fc (orange). Both variants show comparable neutralisation efficiencies (Mean ± SD). Unpaired t-test of AUC (t=1.695, df=28). B) Neutralisation assay of SARS-CoV-1, SARS-CoV-2, SARS-CoV-2 D614G, SARS-CoV-2 B1.1.7 and SARS-CoV-2 B1.351 pseudotyed vectors with ACE2(HH:NN)-Fc (LALA-PG), LY-CoV-55, REGN-COV2 cocktail, REGN10933 and REGN10987. Marked decrease of neutralisation capacity for SARS-CoV-2 B1.351 detected for LY-CoV555, REGN-COV2 cocktail, REGN10933 and REGN10987. No loss of neutralisation capacity detected for ACE2(HH:NN)-Fc (LALA-PG) receptor decoy (Mean ± SD). One way ANOVA of AUC with Dunnett’s multiple comparison to blue (F=369.2, df=4,88). Bottom right panel, fold change of neutralisation capacity based on NT_50_ values. * = unmeasurable NT_50_ value.

Next, to investigate the degree of neutralisation efficiency against the SARS-CoV-2 variants of interest, the receptor decoy was tested against the engineered replication deficient lentiviral vectors pseudotyped with the glycoproteins of SARS-CoV-2 WT, D614G mutation, UK B.1.1.7 and South Africa B.1.351 variants and SARS-CoV-1. The ACE2(HH:NN)-Fc LALA-PG was able to efficiently neutralise SARS-CoV-2, with tight dose-response curves among the SARS-CoV-2 variants, and SARS-CoV-1 (**Figure 5B**). Interestingly, the neutralisation capacity was slightly improved for the B.1.1.7 and B.1.351 variants compared to WT SARS-CoV-2. The monoclonal antibody LY-CoV555 showed a marked reduction in neutralisation capacity for the D614G and B.1.1.7 variants, 3 and 8-fold respectively, significantly impacting on the antibody efficacy; with an almost complete abrogation of neutralisation against the B.1.351 variant (**Figure 5B**). The 1:1 REGN10933/REGN10987 antibody cocktail (REGN-COV2) was more resilient in its response to the SARS-CoV-2 variants but was characterised by a 4-fold reduction in neutralisation for the B.1.351 variant. When the two antibodies constituting the cocktail were analysed individually, the REGN10933 showed a 3-fold decrease in neutralisation capacity for the D614G and B.1.1.7 variants, with a staggering > 1000-fold reduction for the B.1.351 variant; while the REGN10987 showed a 4-fold neutralisation reduction for the B.1.1.7 variant and a 10-fold shift for the D614G variant (**Figure 5B**).

### In vivo neutralisation of SARS-CoV-2 in a hamster model of disease

It has been previously reported that Syrian hamsters *(Mesocricetus auratus)* are a relevant small animal model for SARS-CoV-2 infection, reporting symptoms such as reduced body weight and pathological lesions on the lung^32^. The hamster FcγRs show a different interaction profile with human IgG1 Fc molecules compared to Hamster IgG Fc, nonetheless, the LALA-PG mutation of our construct still showed a complete lack of interaction (**Supplementary Figure 3**). Syrian hamsters were challenged intranasally with 10^4.5^ median tissue culture infectious dose (TCID_50_) viral inoculum and then dosed 24 h later via intra-peritoneal (i.p.) injections of ACE2(HH:NN)-Fc LALA-PG at either 5 mg/kg or 50 mg/kg. PBS injections were used for the placebo control group. The hamster groups treated with either high or low ACE2(HH:NN)-Fc LALA-PG doses showed a significant protection against body weight loss, with a maximum average weight loss of 11% compared to 21% for the placebo group, relative to the day of viral inoculum (**Figure 6A**). Throat swabs revealed a substantial reduction in viral RNA copies between day 4 and day 6 post-viral challenge, compared to the placebo; several animals showed undetectable levels of RNA between day 3 and day 6 for the high ACE2(HH:NN)-Fc LALA-PG dose and an overall viral load significantly lower than placebo control group (**Figure 6B**). Macro-analysis on lung necropsies (day 7) also showed an overall reduction in lung damage for the ACE2(HH:NN)-Fc LALA-PG treated groups, characterised by fewer lesions and blood clotting (**Figure 6C**). Finally, i.p. administered ACE2(HH:NN)-Fc LALA-PG was still detectable in the hamster sera at day 7, with levels almost 20-fold higher for the high dose compared to low dose treatment (**Figure 6D**).

**Figure 6.**
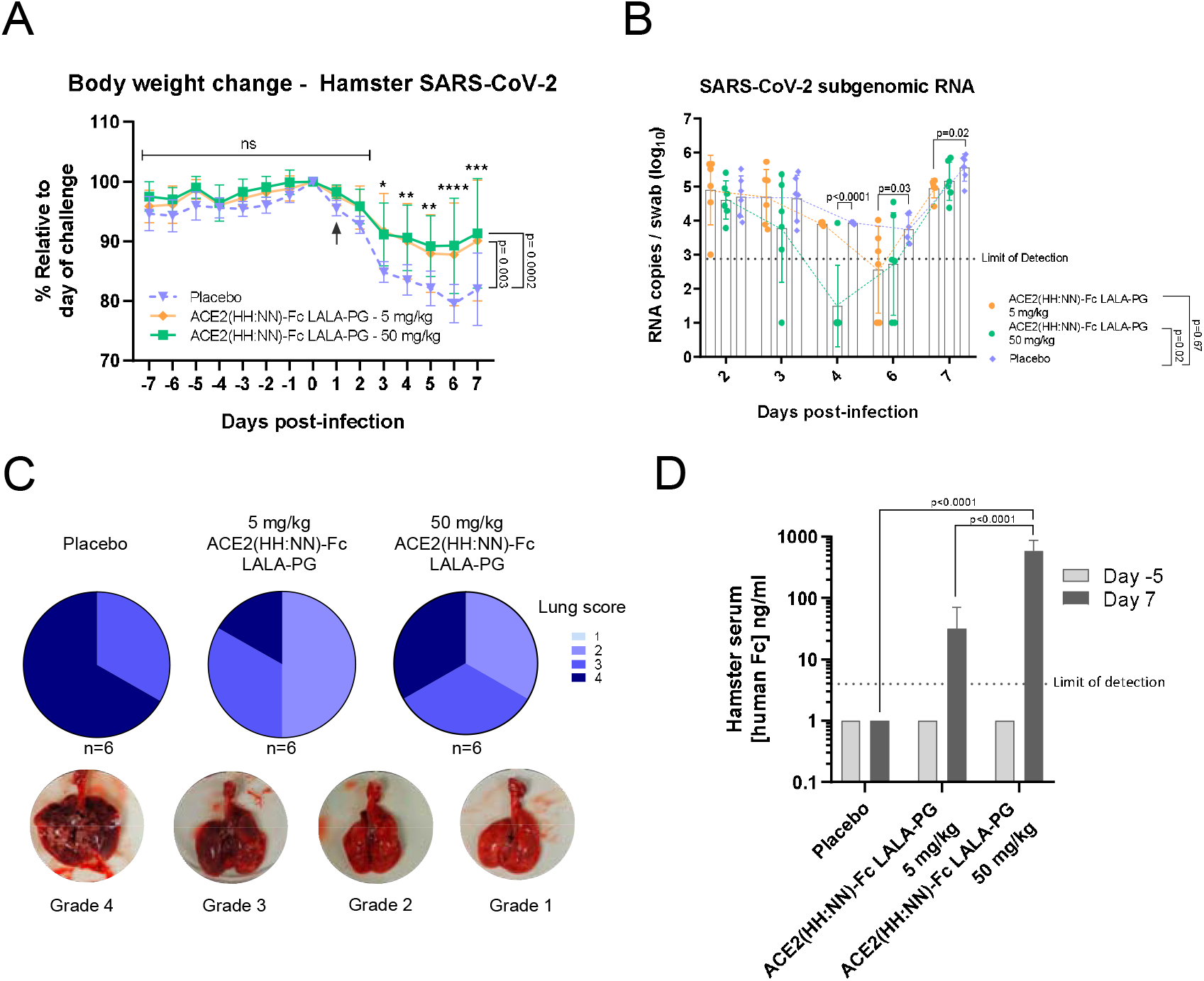
*In vivo* SARS-CoV-2 neutralisation. Syrian hamster intranasally challenged with live SARS-CoV-2. ACE2(HH:NN)-Fc (LALA-PG) administered i.p. at day 1 post-challenge at 5 mg/kg, 50 mg/kg or placebo (PBS) (n=6 per group). A) Body weight change (%) relative to the day of viral inoculation. Day of therapeutic administration marked with arrow. Significant reduction of body weight change relative to placebo, detected for both treatment regimens (Mean ± SD). Individual day comparison placebo vs. 50 mg/kg dose two way ANOVA with Sidak’s multiple comparison compared to placebo group * p=0.01, ** p=0.004, *** p=0.0001, **** p<0.0001. One way ANOVA of AUC with Turkey’s multiple comparison (F=9.379, df=2, 225). B) Sub genomic RNA PCR swab test. Limit of detection 2.88 RNA copies, samples with undetectable RNA were assigned a value of 1 (Mean ± SD). Two way ANOVA with Dunnett’s multiple comparison, one way ANOVA of AUC with Dunnett’s multiple comparison (F=3.247, df=2, 75). C) Necropsy pathology lung score (categories 1-4) showing reduction in lung damage for ACE2(HH:NN)-Fc LALA-PG treated groups. Bottom, representative lung damage for grade score 1, 2 and 3. D) Human IgG1 Fc concentration in hamster sera at day −5 and day 7 relative to viral inoculation. Limit of detection 4 ng/ml. Samples with undetectable levels were assigned a value of 1 (Mean ± SD). Two way ANOVA with Sidak’s multiple comparison (F=39.2, df=2, 22).

### Formulation optimisation and streamlined manufacturing of ACE2(HH:NN)-Fc decoy

To define a suitable formulation considering manufacturing scale-up for clinical application, the well-established antibody formulation buffer 20 mM His^33^, was used to solubilise the ACE2(HH:NN)-Fc at a range of pH conditions from 3.5 to 7. The ACE2(HH:NN)-Fc in PBS at pH 7.4 showed good thermal stability with a first unfolding event at 46.1°C, attributed to the unfolding of the ACE2 domain (**Figure 7A**). When tested in 20 mM His buffer, the first unfolding event occurred at a Tm between 42.3 and 51.6°C, with the lowest Tm associated with pH 3.5 and the most stable Tm obtained at pH 6.5 (**Figure 7A**).

**Figure 7.**
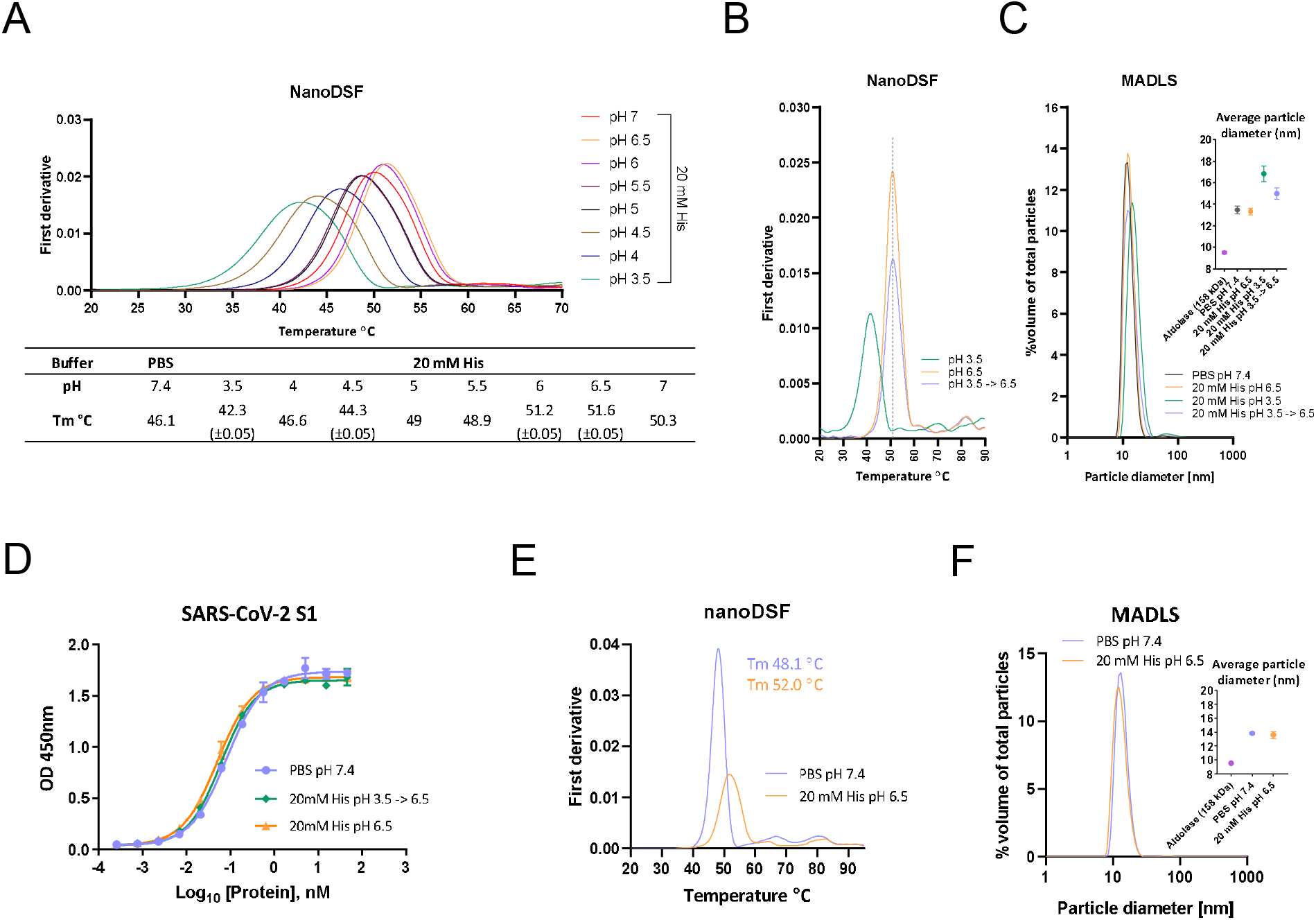
ACE2-Fc formulation optimisation. **A**) Thermal stability analysis via nanoDSF of ACE2(HH:NN)-Fc in PBS at pH 7.4 or in 20mM His buffer with pH range of 3.5-7. Highest stability obtained with 20 mM His pH 6.5. **B**) Thermal stability of ACE2(HH:NN)-Fc in 20 mM His pH 6.5 (orange) and 3.5 (green) following 2h incubation at RT. Full stability could be recovered following buffer exchange of ACE2(HH:NN)-Fc from pH 3.5 to pH 6.5 (blue). **C**) Particle size distribution analysis via MADLS of ACE2(HH:NN)-Fc at 1 mg/ml in PBS pH 7.4, 20 mM His pH3.5, 6.5 or buffer exchanged from pH 3.5 to 6.5. Increase of particle size of sample at pH 3.5 was partially recovered upon buffer exchange in 20 mM His pH 6.5 (Mean ± SD). **D**) ACE2(HH:NN)-Fc binding capacity for SARS-CoV-2 S1 in ELISA in PBS pH 7.4 (blue), 20 mM His pH 6.5 (orange) or 20 mM His pH3.5 followed by buffer exchange in 20 mM His pH 6.5 (green). (Mean ± SD). **E**) Thermal stability analysis via nanoDSF of ACE2(HH:NN)-Fc (LALA-PG) in PBS at pH 7.4 (blue) or in 20 mM His pH 6.5 (orange) showing a 3.9 °C Tm shift. **F**) Particle size distribution analysis via MADLS of ACE2(HH:NN)-Fc (LALA-PG) at 1 mg/ml in PBS pH 7.4 (blue) and 20 mM His pH 6.5 (orange) showing comparable profile (Mean ± SD).

A crucial phase during manufacturing of monoclonal antibodies lies in the viral inactivation step, often carried out at low pH^34^, which can affect the stability and aggregation state of the proteins in solution. To investigate this, the ACE2(HH:NN)-Fc was exposed to pH 3.5 for 90 minutes before dialysis in 20 mM His pH 6.5. Thermal stability comparison of ACE2(HH:NN)-Fc at pH 3.5, 6.5 and 3.5 dialysed to 6.5 showed how the initial instability due to pH 3.5 could efficiently be restored to that of the ACE2(HH:NN)-Fc following dialysis at pH 6.5 (**Figure 7B**). The distribution of particles within the solution showed a predominantly monodispersed profile for the ACE2(HH:NN)-Fc in PBS and 20 mM His pH 6.5, with an average diameter of 13.5 and 13.3 nm, respectively, in agreement with a molecule of predicted MW of 219 kDa. The suspension in a low pH buffer of 3.5 did not significantly enhance aggregation of ACE2(HH:NN)-Fc (**Figure 7C**). Furthermore, the change of buffer from PBS pH 7.4 to 20 mM His pH 6.5 and, crucially, the viral inactivation step in at pH 3.5 with subsequent dialysis to pH 6.5, did not affect the capacity of the ACE2(HH:NN)-Fc to bind the SARS-CoV-2 S1 protein, further validating the proposed process (**Figure 7D**).

The ACE2(HH:NN)-Fc LALA-PG also showed an increased thermal stability when in 20 mM His pH 6.5 buffer, with Tm moving from 48.1°C to 52.0°C, and CH2 CH3 unfolding happening at 64.3°C and 81.8°C, respectively (**Figure 7E**). The ACE2(HH:NN)-Fc LALA-PG was also characterised by a monodispersed particle profile with an average diameter size of 13.6 nm in 20 mM His pH 6.5 (**Figure 7F**). Finally, the formulation in 20 mM His pH 6.5 of ACE2(HH:NN)-Fc LALA-PG did not alter the SARS-CoV-2 neutralisation capacity of the construct (**Supplementary figure 4**).

## DISCUSSION

We have described the generation of a catalytically inactive ACE2 receptor decoy fused to an engineered human Fc domain with abrogated FcγR engagement, showing optimal biophysical properties and manufacturability. The construct showed strong neutralisation potency against several SARS-CoV-2 variants of concern *in vitro* and evidence of efficacy as a therapeutic administration in a live viral challenge model *in vivo.*

Monoclonal antibodies developed for the treatment of COVID-19 have shown efficacy in the treatment of early phases of the infection, potentially useful in prophylaxis or as an alternative for people who cannot be vaccinated^35^. However, cumulative spike-protein mutants may render therapeutic mAbs ineffective. For instance, the variants of concern B.1.351 and P.1 have been shown to affect the neutralisation capacity of the approved antibody therapeutics. The LY-CoV555 antibody reported an almost complete abrogation of neutralisation, while the antibody cocktail REGN-COV2 showed a severe impairment for one of its components, suggesting preservation of limited therapeutic efficacy^17,18^. Although not yet present in naturally occurring variants, the single amino acid mutation E406W has recently been shown to be able to escape both antibodies in the REGN-COV2 cocktail^36^.

“Receptor traps” are an established therapeutic approach, e.g. the anti-TNF Etanercept^19^, the VEGF-trap Aflibercept^20^ and the CTLA-4-Ig Abatacept^21^. Since ACE2, the receptor for SARS-CoV-2, is a type I transmembrane protein with a discrete extracellular domain, ACE2-based receptor decoys may be effective against COVID-19. A theoretical advantage of this approach is resistance to S-protein mutational drift since mutations disrupting interaction with ACE2 would render the virus inactive. ACE2 based therapeutics have been described: A soluble catalytically active human ACE2 showed efficacy in the treatment of a severe COVID-19 patient by reducing plasma viraemia^37^. Two recent reports have described engineered ACE2 molecules with sub nM affinities for the S glycoprotein^23,25^. Similarly, ACE2-derived inhibitory peptides with improved manufacturability and stability, also showed enhanced SARS-CoV-2 neutralisation efficacy^24^. However, the lack of an Fc domain may impact on serum half-life and manufacturing efficiency and importantly optimized designs may allow viral mutational escape due to differences with the endogenous receptor. Plain ACE2-Fc fusions were developed against SARS-CoV-1 in 2003 and was also proposed for SARS-CoV-2^38,39^. Recently, a tetravalent ACE2-Fc was described which showed improved neutralisation efficiency compared to standard ACE2-Fc formats, without ACE2 domain engineering^40^. These ACE2-Fc fusions retained catalytic activity of ACE2 and maintained full Fc effector function.

A key factor in the function of ACE2 based decoys is their affinity to the S-protein and its variants. To our knowledge, comparison of ACE2 affinity and biophysical properties for the SARS-CoV-2 variants S1 domains has not yet been reported. An enhanced affinity has been shown for the RBD domain of the B.1.1.7 and B.1.351 variants, however, this may not reflect the effect of the full S1 domain^41^. To this end, we generated recombinant S1 domains for the SARS-CoV-2 WT (Wuhan Hu-1), D614G, B.1.1.7 and B.1.351 variants to assess ACE2 binding affinities. WT, D614G and B.1.351 S1 domains displayed comparable binding affinity to their receptor, with a slightly slower kinetic profile for the B.1.351. Interestingly, the B.1.1.7 variant showed a > 3-fold enhancement in affinity and we speculate that this feature may have contributed to the rapid spread of the variant. We have also pseudotyped replication deficient lentiviral vectors with either the WT SARS-CoV-2 spike glycoprotein or bearing the D614G, B.1.1.7 and B.1.351 mutations. Interestingly, all pseudotyped vectors exhibited comparable particle productivity, but differed in their infectivity as measured by viral titres on SARS-CoV permissive HEK293T cells. The SARS-CoV-1 pseudotype showed the weakest infectious capacity, 3.2-fold lower than WT SARS-CoV-2, while the D614G variant showed a 2.6-fold increase relative to the parent protein. The B.1.1.7 and B.1.351 pseudotypes showed approximately 1.8-fold increase in infectivity compared to the WT version. These differences may in part be explained by the relative changes in affinity/interaction of the spike proteins for the ACE2 receptor and suggest increased activity of ACE2 based therapeutics against certain variants.

In contrast to previously reported formats, we developed an ACE-Fc decoy engineered to be catalytically inactive to prevent systemic activity, and with a completely abrogated FcγR interaction to minimise pro-inflammatory activity. We provided evidence for lack of enzymatic activity on a synthetic substrate, while showing that these mutations still maintain reversible engagement with the natural substrate Ang II, avoiding the risk of acting as a substrate sink. While the use of an active ACE2 domain may be beneficial to restore the balance in the affected lung, the inactive ACE2 offers a safer profile over systemic interference on the renin/angiotensin system. In contrast to a previously reported ACE2-Fc construct carrying the LALA Fc mutation^42^, our design using the LALA-PG mutation shows a complete abrogation of human FcγR engagement as analysed by SPR and flow cytometry against cell lines and human M1 macrophages, while maintaining FcRn interaction to provide extended half-life^29^. Although its relevance has not been conclusively determined for SARS-CoV-2, the engineered Fc should alleviate the risk of antibody-dependent enhancement (ADE) of infection as reportedly mediated through FcγR II for SARS-CoV-1 and MERS-CoV^43,44^.

Despite the introduced mutations, which sit outside the S-protein targeting region, the ACE2 decoy maintained equivalent affinity and kinetic interactions to those of the active receptor for the SARS-CoV-2 S glycoprotein variants, minimising the risk of mutational escape. Additionally, we showed binding capacity to SARS-CoV-1 and HCoV-NL63, providing evidence for broad-spectrum activity over ACE2-tropic viruses. In a direct comparison, the antibodies LY-CoV555, REGN10933 and REGN10987 showed binding capacity for the SARS-CoV-2 WT, D614G and B.1.1.7 but failed to recognise the S-protein from SARS-CoV-1 and HCoV-NL63. Strikingly, the LY-CoV555 and REGN10933 mAbs were strongly impaired in their binding to the B.1.351 variant. Although the final constant domain sequences used in our version of the aforementioned antibodies may vary compared to the clinical products, the variable domains and antibody formats were generated according to published information^30,31^.

Pseudotyped vectors were used to compare the relative SARS-CoV-2 neutralisation efficiency of the antibodies LY-CoV555, REGN10933, REGN10987 and the 1:1 REGN10933/REGN10987 antibody cocktail REGN-COV2, to the described ACE2-Fc decoy. In line with previous reports^17^, we have observed a substantial drop in neutralisation capacity of the LY-CoV555, the REGN-COV2 cocktail and the latter’s individual antibodies REGN10933 and REGN10987 for the SARS-CoV-2 variants. The LY-CoV555 and REGN10933 were especially impaired by the B.1.351 variant. We also noticed a generally reduced neutralisation capacity for the D614G variant, which is likely due to its enhanced infectious capacity compared to the WT strain. Strikingly, the ACE2 decoy maintained efficacy towards SARS-CoV-1 and all SARS-CoV-2 variants tested, showing enhanced potency driven by mutational drift. Paradoxically, the spike mutations enhancing affinity for the ACE2 receptor would improve the neutralisation potency of ACE2-based decoys, while affecting the performance of mAbs. The ACE2(HH:NN)-Fc LALA-PG was also able to affect the replication of SARS-CoV-2 live virus in a hamster model, reducing body weight loss and lung damage in infected animals.

As a fusion construct, unexpected off-target binding events could manifest, representing a liability over its safety profile. To this end, a cross-reactivity study against a comprehensive panel of close to 6000 human soluble and membrane-bound proteins has highlighted the exquisite specificity of this construct for the target protein, providing confidence over its safety profile.

Finally, we achieved high expression yields for our ACE2 receptor decoy, using a non-optimised transient transfection system and single-step protein-A affinity purification. The stability of the construct at pH 3.5, while showing no change in its aggregation profile and binding capacity, lends to its suitability for antibody-like purification processes such as low pH viral inactivation^45^. We have also determined an optimal pH to enhance stability of the protein using a commonly adopted His buffer for clinical-stage monoclonal antibodies^46^, raising the Tm to 52°C. These characteristics would allow for standard manufacturing scale-up required for clinical grade material.

In conclusion, we describe detailed *in vitro* and *in vivo* characterisation of a soluble catalytically inactive ACE2-Fc receptor decoy molecule resistant to spike protein mutation. We also demonstrate that our decoy molecule has the potential for rapid upscale manufacturability. In theory, our decoy should be active against any new ACE2-tropic virus which might emerge in the future. In this phase of the SARS-CoV-2 pandemic where viral variants are exerting pressure over the efficacy of vaccines and monoclonal antibodies, the development of biotherapeutics which are inherently resistant to SARS-CoV-2 mutations may be prudent.

## MATERIALS and METHODS

### Cell line generation and maintenance

HEK-293T (ATCC – CRL-11268) were cultured in Iscove’s modified Dulbecco’s medium (IMDM) (Lonza – 12-726F) supplemented with 10% Foetal Calf Serum (FCS, Biosera – FB 1001/500) and 2 mM GlutaMAX™ (Gibco – 35050061) at 37°C with 5% CO_2_. Sup-T1 (ATCC – CRL-1942), U937 (ATCC – CRL-1593.2) and K562 (ATCC – CCL-243) were cultured in Roswell Park Memorial Institute (RPMI) 1640 medium (Gibco – 21875034) supplemented with 10% Foetal Calf Serum (FCS, Biosera – FB 1001/500) and 2 mM GlutaMAX™ (Gibco – 35050061) at 37°C with 5% CO_2_.

Sup-T1 cells were γ-retrovirally transduced to express the S glycoprotein of SARS-CoV-2 Wuhan Hu-1 strain co-expressed with eBFP as a marker gene. Briefly, 5 x 10^5^ cells were incubated with 1mL of unquantified vector supernatant in the presence of retronectin in non-TC treated 24-well plate and subjected to spin-inoculation at 1000 g for 40 mins. Cells were recovered 24 h later by culturing in serum supplemented RPMI 1640 for two passage before use in experiments.

For the generation of MDM, human monocytes were isolated from blood of healthy donors using Easy Sep human monocyte isolation kit (Stemcell – 19359), according to manufacturer’s recommendations. Monocyte isolation was determined with the following flow cytometry antibody panel after 10 min incubation with anti-human CD32 (StemCell – 18520): APC anti-human CD14 (Biolegend – 301808), PE-Cy7 anti-human CD3 (Biolegend – 344186), AF488 anti-human CD20 (Biolegend – 302316) and live/dead Sytox Blue stain (Invitrogen – S34857). Monocytes were activated by culturing in Immunocult SF macrophage differentiation media (Stemcell – 10961) supplemented with 50 ng/ml M-CSF (Stemcell – 78057). At day 6, cells were supplemented with 50 ng/ml IFN-γ (Stemcell – 78020) and 10 ng/ml LPS (Sigma – L4391) to stimulate M1 polarization. M1 macrophages were harvested by Accutase dissociation (Stemcell – 07920). The following flow cytometry antibody panel was used to determine monocyte differentiation and M1 polarization after 10 min incubation with anti-human CD32: APC anti-human CD14 (Biolegend – 344186) BV421 anti-human CD80 (Biolegend – 305222), PE anti-human CCR7 (Biolegend 353204), APC/Fire750 anti-human CD209 (330116) and 7-AAD viability staining solution at 5 μl/1×10^6^ cells. Samples from both flow staining panels were acquired using the MacsQuant10 instrument (Miltenyi Biotech).

### Protein expression, purification, and characterisation

Human ACE2 amino acid 18-740 (Uniprot Q9BYF1) was fused to the human IgG1 hinge and Fc (Uniprot P01857). Inactive ACE2 was generated by introducing H374N and H378N mutations. Silent Fc variants were generated with L234A/L235A and L234A/L235A/P329G mutations. Chimeric human inactive ACE2 with hamster Fc fusion was generated using Cricetulus migratorius IgG heavy chain hinge-Fc sequence (GenBank U17166.1). Variable domain sequences for LY-CoV-555 was obtained from published crystal structure (PBD 7L3N)^30^, REGN10933 and REGN10987 sequences were obtained from published crystal structure (PDB 6XDG)^31^. Heavy variable domains were fused to human IgG1 constant chain (Uniprot P01857); kappa variables were fused to human kappa constant domain (Uniprot P01834); lambda variable was fused to human lambda constant 1 (Uniprot P0CG04). All constructs were cloned in an AbVec vector^47^. REGN-COV2 antibody cocktail was generated as a 1:1 mix of REGN10933 and REGN10987. Recombinant Fc tagged proteins were expressed by transient transfection in ExpiCHO, according to manufacturer’s recommendations (Thermo Fisher – A29133). Supernatant from transfected CHO cells was purified using 1 ml HiTrap MabSelect PrismA (GE Healthcare – 17549851) affinity chromatography with in-line dialysis in PBS via HiTrap 5 ml desalting columns (GE Healthcare – 29048684) using an Akta™ Pure system (GE Healthcare), following manufacturer’s recommendations.

SARS-CoV-2 S1 domains (aa 1-681) from WT (GenBank – QHD43416.1) or including the D614G^11^, B.1.1.7^13^ (aa 1-680) and B.1.351^14^ mutations, were cloned in fusion with a dual 6xHis tag using an AbVec vector. Supernatant from Expi293 transfected cells was manually purified using TALON metal affinity chromatography (Takara bio Inc – 635502), according to manufacturer’s recommendations. Purified proteins were buffer exchanged in PBS using Zeba spin desalting columns (Thermo Fisher – 89890).

Purified proteins were analysed for purity determination via sodium dodecyl sulfate polyacrylamide gel electrophoresis (SDS-PAGE) on a 4-20% gradient gel (BioRad – 4568094), with or without the presence of 2-mercaptoethanol as reducing agent.

### Differential scanning fluorimetry

Thermal stability was determined by differential scanning fluorimetry nano(DSF) on a Prometheus NT.48 instrument (Nanotemper) using first derivative of 350/330nm ratio to determine the melting temperature (Tm) value. Samples were loaded on a glass capillary and temperature scanned from 20 to 95 °C at 1°C/min.

### Aggregation and particle size measurement

Aggregation propensity and average particle size of the test proteins was determined using a Zetasizer Ultra device and ZS Xplorer software (Malvern Panalytical) by MADLS. Samples were loaded into a low volume quartz cuvette (Malvern Panalytical – ZEN2112) at a concentration of 1 mg/ml or 20 mg/ml. Triplicate measurements were taken for each sample. Particle size of Aldolase (158 kDa) was used as reference.

### ACE2 enzymatic activity

Enzymatic activity of active ACE2-Fc (ACRO biosystems – AC2-H5257) and ACE2(HH:NN)-Fc was measured by using Mca-APK(Dnp) (Enzo Life Science – BML-P163) as substrate in 96-well black microtiter plates. Samples were diluted in reaction buffer (50 mM 4-morpholineethanesulfonic acid, pH = 6.5, 300 mM NaCl, 10 μM ZnCl_2_ and 0.01% Triton X-100) at a concentration of 0.1 μg/ml in the presence of 20 μM of Ma-APK(Dnp) or control peptide BML-P127 (Enzo Life Sciences) in a final volume of 100 μl/well. The reaction was performed in triplicate at room temperature for 1h. Activity was measured as fluorescence intensity at 320 nm/393 nm (Ex/Em) wavelength at 1-minute intervals using a Varioskan LUX instrument (Thermo Scientific).

### ELISA on spike protein

Nunc Maxisorp clear 96-well plates were coated with 1 μg/ml (in PBS) of SARS-CoV-2 S trimer (ACRO biosystems – SPN-C52H9), SARS-CoV-2 S1 domain (ACRO biosystems – S1N-C52H3) or BSA (Sigma – A7906) overnight at 4°C in 50 μl/well. Plates were blocked with PBS 2% BSA for 1h at RT. Test proteins were incubated at 45.6 nM concentration with 3-fold serial dilutions for 1h at RT in PBS 0.5% BSA. Bound Fc-tagged proteins were detected with anti-human HRP-conjugated secondary antibodies (Jackson ImmunoResearch – 109-035-088) at 1:3000 dilution in PBS 0.5% BSA. Incubation was allowed for 1h at RT. All washes were performed in PBS 0.05% Tween20. Specific interaction revealed with 1- step TMB Ultra reagent (Thermo Fisher – 34028) at 45 μl/well and blocked with 45 μl/well of 1M H_2_SO_4_. Plates were acquired on a Varioskan Lux instrument at a wavelength of 450 nm. Data analysed with GraphPad Prism 8 (GraphPad software).

### Flow cytometry

For FcγR binding assay on U937, K562, SupT1 cells and MDM, test constructs were incubated at specified concentrations for 30 minutes at 4°C to prevent dissociation/internalisation. Protein labelled cells were stained with biotinylated SARS-CoV-2 S1 (ACRO biosystems – S1N-C82E8) and detected with streptavidin AF647 (Invitrogen – S21374). Cells were stained with 7-AAD viability staining solution at 5 μl/1×10^6^ cells to determine live cells. Stained samples were acquired using a MacsQuant10 instrument (Miltenyi Biotec) and analyzed on FlowJo software (BD).

Binding capacity of ACE2(HH:NN) Fc, LALA Fc and LALA-PG Fc to SupT1 expressing wild-type SARS-CoV-2 full length spike was assessed via incubation of test protein at 45.6 nM with 2-fold serial dilutions for 30 mins at RT, followed by secondary incubation with anti-Human IgG (H+L) AF647 (Invitrogen – A21445) for 20 mins at RT in the dark. Cells were stained with 7-AAD viability staining solution at 5 μl/1×10^6^ cells to determine live cells and subsequently acquired using MacsQuant10 instrument. Flow cytometry data was analyzed on FlowJo software (BD).

### Surface plasmon resonance

Recombinant active ACE2-Fc (ACRO biosystems – AC2-H5257) and ACE2(HH:NN)-Fc constructs were captured on flow cell 2 of a Series S Protein A sensor chip (GE Healthcare – 29127555) to a density of 500 RU using a Biacore 8K instrument (GE Healthcare). HBS-EP^+^ buffer was used as running buffer in all experimental conditions. Recombinant purified Angiotensin II (Sigma – A9525) at 1 μM with 2-fold serial dilutions, was used as the ‘analyte’ and injected over the flow channels with 150s contact time and 500s dissociation.

For SARS-CoV-1 S1 (ACRO biosystems – S1N-S52H5), HCoV-NL63 S1 (SIN-V52H3), SARS-CoV-2 S1 WT (ACRO biosystems – S1N-C52H3) and in-house expressed SARS-CoV-2 S1 WT, D614G, B.1.1.7 and B. 1.351 kinetics, test ACE2-Fc constructs and antibodies were captured to a density of 70 RU or 50 RU, respectively, on a Series S Protein A sensor chip (GE Healthcare – 29127555) using a Biacore T200 and Biacore 8k instruments (GE Healthcare). HBS-P+ buffer was used as running buffer in all experimental conditions. Recombinant purified spike proteins at known concentrations were used as the ‘analyte’ and injected over the respective flow cells with 150s contact time and 300s dissociation.

The binding kinetics to FcγRIa (ACRO biosystems – FCA-H52H1), FcγRIIa (ACRO biosystems – CD1- H5223), FcγRIIb (ACRO biosystems – CDB-H5228), FcγRIIIa (ACRO biosystems – CDA-H5220) and FcγRIIIb (ACRO biosystems – CDB-H5222) were captured to a density of 50 RU (or 150 RU for FcγRIIa, FcγRIIb and FcγRIIIb) on flow cell 2, 3 or 4 of a Series S CM5 chip (GE Healthcare) functionalised with an anti-His capture kit (GE Healthcare) using a Biacore T200 instrument. HBS-EP+ buffer was used as running buffer in all experimental conditions. Purified ACE2(HH:NN)-Fc, ACE2(HH:NN)-Fc LALA and ACE2(HH:NN)-Fc LALA-PG at a concentration of 500 nM with 2-fold serial dilutions were used as the ‘analyte’ and injected over the respective flow cells with 150s contact time and 300s dissociation.

All experiments were performed at 25°C with a flow rate of 30 μl/ml. Flow cell 1 was unmodified and used for reference subtraction. A ‘0 concentration’ sensogram of buffer alone was used as a double reference subtraction to factor for drift. Data were fit to a 1:1 Langmuir binding model using Biacore insight evaluation software (GE Healthcare). SARS-CoV-1 S1 sensograms were also fit to a two-state kinetics. Since a capture system was used, a local Rmax parameter was used for the data fitting in each case.

### Viral vector production

Viral vectors were produced by triple transient transfection of HEK-293Ts in 100 mm plates using GeneJuice (Merck – 70967) with a total of 12.5 μg of DNA. γ-retroviral vectors were produced by triple transient transfection of 4.69 μg Peq-Pam plasmid (encoding Moloney GagPol), 3.13 μg of RDF plasmid (encoding RD114 envelope) and 4.69 μg retroviral backbone SFG^48^ expressing full-length SARS-CoV-2 S glycoprotein co-expressed with eBFP as marker gene. Similarly, for lentiviral vector production, cells were transfected with 5.42 μg of pCMV-dR8.74 (encoding lentiviral GagPol), 2.92 μg of envelope plasmid expressing codon-optimised SARS-CoV S glycoproteins with their ER retention signals deleted (deletion of the last 19 amino acids on the carboxy-terminus) and 4.17 μg of lentiviral backbone pCCL encoding eGFP as transgene driven by internal viral SFFV promoter.

Culturing medium was changed 24 h post-transfection and vector supernatants were collected 48 h after transfection and processed by centrifugation at 1000 g for 10 mins at 4°C to remove cellular debris followed by microfiltration using Millex-HV 0.45 μm syringe filter units (Merck – SLHV033RB). Viral supernatants were either kept on ice for further use or frozen down at −80°C for storage.

### p24 ELISA

Physical particles were determined by measuring p24 levels using the QuickTitre™ Lentivirus Titre which quantifies lentivirus-associated HIV rather than free p24 proteins (Cell Biolabs – VPK-107-T). Manufacturer’s protocol was followed, and samples were assayed in triplicates. Briefly, after incubation with kit’s ViraBind™ reagents and virus inactivation, samples were incubated in microwell plates precoated with anti-p24 antibodies followed by a subsequent incubation with secondary FITC-conjugated anti-HIV p24 monoclonal antibody (1:1000). Subsequently, well were exposed to HRP-conjugated anti-FITC monoclonal antibody (1:1000). Plates were acquired on a Varioskan Lux instrument at a wavelength of 450 nm. Data analysed with Graph Prism 8 (GraphPad software).

### SARS-CoV-2 lentiviral pseudotyped viral vector titration

Functional infectious viral titres were determined by flow cytometry analysis (BD LSRFORTESSA X-20 cell analyser) of transgene expression in transduced HEK-293T cells that were previously engineered to express human ACE2 and TMPRSS2. Experiments were performed in 24-well plates (50,000 cells/well). Serially diluted viral supernatants were added onto seeded cells in the presence of 8 μg/mL polybrene. Transduction efficiencies were determined 72 h later using BD LSRFORTESSA X-20 cell analyser and eGFP expression between 0.5% – 20% were used in the following equation to determine viral titer:

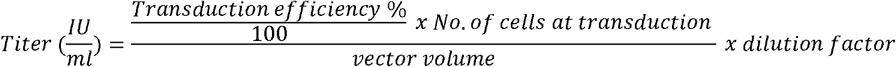

### SARS-CoV-2 lentiviral pseudotyped viral vector neutralisation assay

Proteins was serially diluted in PBS to 7 decreasing concentrations ranging from 100 mg/mL to 6.1 ng/mL (4-fold serial dilution). Each antibody dilution was then mixed 1:1 with lentiviral vectors pseudotyped with SARS-CoV S glycoproteins normalised to 1.0 x 10^5^ physical particle of vectors pseudotyped with WT Wuhan Hu-1 glycoprotein, to a final volume of 200 μL and incubated at 37 °C for 1 h. Antibody-virus mixtures were then cultured with 3 x 10^4^ HEK-293T cells previously genetically engineered to express human ACE2 and TMPRSS2, in the presence of 8 μg/mL of polybrene, in 48-well plates with a final volume of 0.5 mL per well. Plates were spin-inoculated at 1000 *g* for 10 mins and incubated for 72 h. Viral titers were then quantified by eGFP expression in target cells using BD LSRFORTESSA X-20 cell analyser and infectivity of all fractions was determined as a percentage of viral titers in the PBS only control.

### SARS-CoV-2 virus neutralisation assay

Vero cells (ATC-CCL81) cultured in Dulbecco’s MEM (Sigma, Cat. No. D6546) with 10% FCS and 2 mM L-Glutamine (Sigma, Cat No. G7513) and 1% penicillin/streptomycin (Invitrogen cat no. 15140148) were seeded the day prior to infection at 2 x10^4^ cells per well in 96-well flat bottom plate. Serial dilutions of proteins of interest were incubated with 100 TCID_50_ of SARS-CoV-2 (strain England/02/2020) for 1 h at 37 °C, 5% CO_2_. After careful removal of culturing media from Vero monolayer, 100μL of protein-virus mixtures were added to the cells and incubated 1 h at 37 °C, 5% CO_2_. Subsequently, 100μL of culturing medium with 4% FBS was added to occupied wells and plates were incubated at 37°C, 5% CO_2_ for 48 h. After removal of culturing medium, occupied wells were fixed with 4% PFA in PBS for 1 h at room temperature for viral inactivation, followed by incubation with 0.1% Triton-X100 for 15 min at room temperature for cell permeabilisation. Plates were washed with 0.05% v/v PBS-Tween and sequentially incubated with mouse anti-SARS-CoV-2 N protein antibody (The Native Antigen Company – MAB12183-100) at 1:500 dilution and HRP-conjugated goat anti-mouse IgG antibody (Jackson ImmunoResearch – 115-035-146) at 1:5000 dilution in 3% w/v milk in 0.05%PBS-Tween. Plates were acquired on a BMG Fluostar Omega at a wavelength of 450 nm.

### Cell microarray test

5477 expression vectors, encoding both ZsGreen1 and a full-length human plasma membrane protein or a cell surface-tethered human secreted protein, and 371 human heterodimers were co-arrayed across a microarray slide in duplicate (**Supplementary Table 1).** HEK293 cells were used for reverse transfection and expression. Test protein was incubated at 20 μg/ml upon cell fixation. Hit detected by fluorescent secondary antibody using ImageQuant software (GE Healthcare). An expression vector (plRES-hEGFR- IRES-ZsGreen1) was spotted in quadruplicate on every slide and used as transfection control. Assay performed by Retrogenix Ltd.

### In vivo hamster model

Syrian hamsters *(Mesocricetus auratus)* RjHan:AURA strain, male and females 4-10 weeks old, were individually caged in a human biosafety level 3 laboratory. At day 0, animals were challenged by intranasal inoculum of 0.1 ml of SARS-CoV-2 with a dose of 10^4.5^ TCID_50_ under medetomidine and ketamine sedation. At day post-inoculum (DPI) 1, animals were treated with i.p. injections of 5 mg/kg or 50 mg/kg ACE2(HH:NN)-Fc LALA-PG or PBS, at equal volumes, and monitored until DPI 7 (group n=6). Non-terminal blood samples were collected at DPI −5 and terminal bleed at DPI 7. For human Fc detection, blood samples were inactivated by incubation at 56 °C for 2h before storing at −20 °C. Detection of residual human Fc in the hamster sera was performed by ELISA, using an anti-human Fc mAb (Sigma-Aldrich, I6260) as capture. The standard curve was generated using purified ACE2(HH:NN)-Fc LALA-PG, passed through the same heat-inactivation step as the serum samples and ranging from 22.8 nM to 22.3 pM via a 2-fold serial dilution. Briefly, Nunc Maxisorp clear 96-well plates were coated with 1 μg/ml (in PBS) of anti-human Fc mAb overnight at 4 °C in 50 μl/well. Plates were blocked the following morning and subsequently loaded with the standards as well as with the serum samples, added at both 1:100 and 1:1000 dilutions. Detection, measurements, and data analysis were performed according to the ELISA protocol described above.

Body weight measurements were recorded daily throughout the study. Throat swabs were collected at DPI 2, 3, 4, 6 and 7 to test for presence of SARS-CoV-2 by qPCR. RNA was isolated using a Direct-zol RNA Miniprep kit (Zymo Research – R2056) and sub genomic RNA detected as previously described^49,50^. Post-mortem examinations were performed at DPI 7. Macroscopic lung lesions were assessed by a pathologist according to the following scoring scheme: 0 = no macroscopical changes; 1 = focal discoloration of lung < 20%; 2 = multifocal discoloration of lung 20-50%; 3 = multifocal discoloration of lung 50-80%; 4 = whole lung affected > 80%. Animal caretakers and pathology personnel were blinded for the treatment groups. Experiments performed by Wageningen Bioveterinary Research Division Virology of Wageningen University. Animal work approved by the Dutch Central Authority for Scientific procedures on Animal (CCD), experimental application 2020.D-0007.016 by the Animal Welfare Body of Wageningen University and Research.

### Statistical analysis

All statistical analyses were performed using GraphPad Prism 8 (GraphPad Software). Specific analysis is detailed in figure legends. A p value < 0.05 was considered significant.

## ACKNOWLEDGEMENTS

This work was funded by the UK Research and Innovation business-led innovation in response to global disruption grant. England/02/2020 isolate of SARS-CoV-2 was kindly provided by Public Health England (PHE), UK. The authors would like to thank Dr Nadia Oreshkova for helpful discussions and guidance in the *in vivo* assays.

## CONTRIBUTIONS

M.F., S.C.O. and M.P. designed the study. M.F., S.C.O., F.T.I. and R.B. planned and/or performed protein purification and biophysical characterisations. J.S., K.L., K.W., F.P., C.A., planned and/or performed plasmid design and cloning. L.M., Z.A. and R.K., planned and/or performed lentiviral production and characterisation and flow cytometry assays. Z.A., and V.B., planned and performed MDM work. L.M. J.S. and P.W. planned and performed cell line development. G.M., E.M.B. and Y.T. planned and/or performed live virus neutralisation assays. A.K., M.F. S.C.O., planned *in vivo* and cross-reactivity studies. M.F. wrote the paper and all authors reviewed the manuscript.

## Notes

### Competing Interest Statement

MF, LM, FTI, ZA, RB, KL, KW, FP, RK, CA, PW, VB, JS, PD, AK, MP and SCO are employees of Autolus Therapeutics.

## REFERENCES

1. Zhou, P. et al. A pneumonia outbreak associated with a new coronavirus of probable bat origin. Nature 579, 270–273 (2020).

2. Huang, C. et al. Clinical features of patients infected with 2019 novel coronavirus in Wuhan, China. The Lancet 395, 497–506 (2020).

3. Wang, D. et al. Clinical Characteristics of 138 Hospitalized Patients With 2019 Novel Coronavirus-Infected Pneumonia in Wuhan, China. JAMA 323, 1061 (2020).

4. Schett, G., Sticherling, M. & Neurath, M. F. COVID-19: risk for cytokine targeting in chronic inflammatory diseases? Nat. Rev. Immunol. 20, 271–272 (2020).

5. Rota, P. A. Characterization of a Novel Coronavirus Associated with Severe Acute Respiratory Syndrome. Science 300, 1394–1399 (2003).

6. Hoffmann, M. et al. SARS-CoV-2 Cell Entry Depends on ACE2 and TMPRSS2 and Is Blocked by a Clinically Proven Protease Inhibitor. Cell (2020) doi:10.1016/j.cell.2020.02.052.

7. Peacock, T. P. et al. The furin cleavage site of SARS-CoV-2 spike protein is a key determinant for transmission due to enhanced replication in airway cells. bioRxiv 2020.09.30.318311 (2020) doi:10.1101/2020.09.30.318311.

8. Raybould, M. I. J., Kovaltsuk, A., Marks, C. & Deane, C. M. CoV-AbDab: the Coronavirus Antibody Database. Bioinformatics btaa739 (2020) doi:10.1093/bioinformatics/btaa739.

9. Yang, L. et al. COVID-19 antibody therapeutics tracker: a global online database of antibody therapeutics for the prevention and treatment of COVID-19. Antib. Ther. 3, 205–212 (2020).

10. Yang, J. et al. A vaccine targeting the RBD of the S protein of SARS-CoV-2 induces protective immunity. Nature 586, 572–577 (2020).

11. Korber, B. et al. Tracking Changes in SARS-CoV-2 Spike: Evidence that D614G Increases Infectivity of the COVID-19 Virus. Cell S0092867420308205 (2020) doi:10.1016/j.cell.2020.06.043.

12. Hodcroft, E. B. et al. Emergence and spread of a SARS-CoV-2 variant through Europe in the summer of 2020. http://medrxiv.org/lookup/doi/10.1101/2020.10.25.20219063 (2020) doi:10.1101/2020.10.25.20219063.

13. Wise, J. Covid-19: New coronavirus variant is identified in UK. BMJ m4857 (2020) doi:10.1136/bmj.m4857.

14. Tegally, H. et al. Emergence and rapid spread of a new severe acute respiratory syndrome-related coronavirus 2 (SARS-CoV-2) lineage with multiple spike mutations in South Africa. http://medrxiv.org/lookup/doi/10.1101/2020.12.21.20248640 (2020) doi:10.1101/2020.12.21.20248640.

15. Greaney, A. J. et al. Comprehensive mapping of mutations to the SARS-CoV-2 receptor-binding domain that affect recognition by polyclonal human serum antibodies. http://biorxiv.org/lookup/doi/10.1101/2020.12.31.425021 (2021) doi:10.1101/2020.12.31.425021.

16. Felipe Naveca et al. Phylogenetic relationship of SARS-CoV-2 sequences from Amazonas with emerging Brazilian variants harboring mutations E484K and N501Y in the Spike protein. https://virological.org/t/phylogenetic-relationship-of-sars-cov-2-sequences-from-amazonas-with-emerging-brazilian-variants-harboring-mutations-e484k-and-n501y-in-the-spike-protein/585 (2021).

17. Wang, P. et al. Antibody Resistance of SARS-CoV-2 Variants B.1.351 and B.1.1.7. http://biorxiv.org/lookup/doi/10.1101/2021.01.25.428137 (2021) doi:10.1101/2021.01.25.428137.

18. Wang, P. et al. Increased Resistance of SARS-CoV-2 Variant P.1 to Antibody Neutralization. http://biorxiv.org/lookup/doi/10.1101/2021.03.01.433466 (2021) doi:10.1101/2021.03.01.433466.

19. Weinblatt, M. E. et al. A Trial of Etanercept, a Recombinant Tumor Necrosis Factor Receptor:Fc Fusion Protein, in Patients with Rheumatoid Arthritis Receiving Methotrexate. N. Engl. J. Med. 340, 253–259 (1999).

20. Holash, J. et al. VEGF-Trap: A VEGF blocker with potent antitumor effects. Proc. Natl. Acad. Sci. 99, 11393–11398 (2002).

21. Genovese, M. C. et al. Abatacept for Rheumatoid Arthritis Refractory to Tumor Necrosis Factor α Inhibition. N. Engl. J. Med. 353, 1114–1123 (2005).

22. Khan, A. et al. A pilot clinical trial of recombinant human angiotensin-converting enzyme 2 in acute respiratory distress syndrome. Crit. Care 21, 234 (2017).

23. Glasgow, A. et al. Engineered ACE2 receptor traps potently neutralize SARS-CoV-2. Proc. Natl. Acad. Sci. 117, 28046–28055 (2020).

24. Linsky, T. W. et al. De novo design of potent and resilient hACE2 decoys to neutralize SARS-CoV-2. Science 370, 1208–1214 (2020).

25. Chan, K. K. et al. Engineering human ACE2 to optimize binding to the spike protein of SARS coronavirus 2. Science 369, 1261–1265 (2020).

26. Schlothauer, T. et al. Novel human IgG1 and IgG4 Fc-engineered antibodies with completely abolished immune effector functions. Protein Eng. Des. Sel. 29, 457–466 (2016).

27. Manson, J. J. et al. COVID-19-associated hyperinflammation and escalation of patient care: a retrospective longitudinal cohort study. Lancet Rheumatol. 2, e594–e602 (2020).

28. Hezareh, M., Hessell, A. J., Jensen, R. C., van de Winkel, J. G. J. & Parren, P. W. H. I. Effector Function Activities of a Panel of Mutants of a Broadly Neutralizing Antibody against Human Immunodeficiency Virus Type 1. J. Virol. 75, 12161–12168 (2001).

29. Lo, M. et al. Effector-attenuating Substitutions That Maintain Antibody Stability and Reduce Toxicity in Mice. J. Biol. Chem. 292, 3900–3908 (2017).

30. Jones, B. E. et al. LY-CoV555, a rapidly isolated potent neutralizing antibody, provides protection in a non-human primate model of SARS-CoV-2 infection. http://biorxiv.org/lookup/doi/10.1101/2020.09.30.318972 (2020) doi:10.1101/2020.09.30.318972.

31. Hansen, J. et al. Studies in humanized mice and convalescent humans yield a SARS-CoV-2 antibody cocktail. Science 369, 1010–1014 (2020).

32. Imai, M. et al. Syrian hamsters as a small animal model for SARS-CoV-2 infection and countermeasure development. Proc. Natl. Acad. Sci. 202009799 (2020) doi:10.1073/pnas.2009799117.

33. Baek, Y., Singh, N., Arunkumar, A. & Zydney, A. L. Effects of Histidine and Sucrose on the Biophysical Properties of a Monoclonal Antibody. Pharm. Res. 34, 629–639 (2017).

34. Mattila, J. et al. Retrospective Evaluation of Low-pH Viral Inactivation and Viral Filtration Data from a Multiple Company Collaboration. PDA J. Pharm. Sci. Technol. 70, 293–299 (2016).

35. Cohen, M. S. Monoclonal Antibodies to Disrupt Progression of Early Covid-19 Infection. N. Engl. J. Med. 384, 289–291 (2021).

36. Starr, T. N. et al. Prospective mapping of viral mutations that escape antibodies used to treat COVID-19. Science 371, 850–854 (2021).

37. Zoufaly, A. et al. Human recombinant soluble ACE2 in severe COVID-19. Lancet Respir. Med. 8, 1154–1158 (2020).

38. Moore, M. J. et al. Retroviruses Pseudotyped with the Severe Acute Respiratory Syndrome Coronavirus Spike Protein Efficiently Infect Cells Expressing Angiotensin-Converting Enzyme 2. J. Virol. 78, 10628–10635 (2004).

39. Lei, C. et al. Potent neutralization of 2019 novel coronavirus by recombinant ACE2-Ig. bioRxiv 2020.02.01.929976 (2020) doi:10.1101/2020.02.01.929976.

40. Miller, A. et al. A Super-Potent Tetramerized ACE2 Protein Displays Enhanced Neutralization of SARS-CoV-2 Virus Infection. https://www.researchsquare.com/article/rs-151560/v1 (2021) doi:10.21203/rs.3.rs-151560/v1.

41. Ramanathan, M., Ferguson, I. D., Miao, W. & Khavari, P. A. SARS-CoV-2 B.1.1.7 and B.1.351 Spike variants bind human ACE2 with increased affinity. http://biorxiv.org/lookup/doi/10.1101/2021.02.22.432359 (2021) doi:10.1101/2021.02.22.432359.

42. Iwanaga, N. et al. Novel ACE2-IgG1 fusions with improved in vitro and in vivo activity against SARS-CoV2. http://biorxiv.org/lookup/doi/10.1101/2020.06.15.152157 (2020) doi:10.1101/2020.06.15.152157.

43. Jaume, M. et al. Anti-Severe Acute Respiratory Syndrome Coronavirus Spike Antibodies Trigger Infection of Human Immune Cells via a pH- and Cysteine Protease-Independent Fc R Pathway. J. Virol. 85, 10582–10597 (2011).

44. Wang, S.-F. et al. Antibody-dependent SARS coronavirus infection is mediated by antibodies against spike proteins. Biochem. Biophys. Res. Commun. 451, 208–214 (2014).

45. Wälchli, R. et al. Understanding mAb aggregation during low pH viral inactivation and subsequent neutralization. Biotechnol. Bioeng. 117, 687–700 (2020).

46. Warne, N. W. Development of high concentration protein biopharmaceuticals: The use of platform approaches in formulation development. Eur. J. Pharm. Biopharm. 78, 208–212 (2011).

47. Tiller, T. et al. Efficient generation of monoclonal antibodies from single human B cells by single cell RT-PCR and expression vector cloning. J. Immunol. Methods 329, 112–124 (2008).

48. Büeler, H. & Mulligan, R. C. Induction of antigen-specific tumor immunity by genetic and cellular vaccines against MAGE: enhanced tumor protection by coexpression of granulocyte-macrophage colony-stimulating factor and B7-1. Mol. Med. Camb. Mass 2, 545–555 (1996).

49. Corman, V. M. et al. Detection of 2019 novel coronavirus (2019-nCoV) by real-time RT-PCR. Eurosurveillance 25, (2020).

50. Wölfel, R. et al. Virological assessment of hospitalized patients with COVID-2019. Nature 581, 465–469 (2020).

