## Supplementary figures for "Characterisation of a novel ACE2-based therapeutic with enhanced rather than reduced activity against SARS-CoV2 variants"

SUPPLEMENTARY MATERIAL

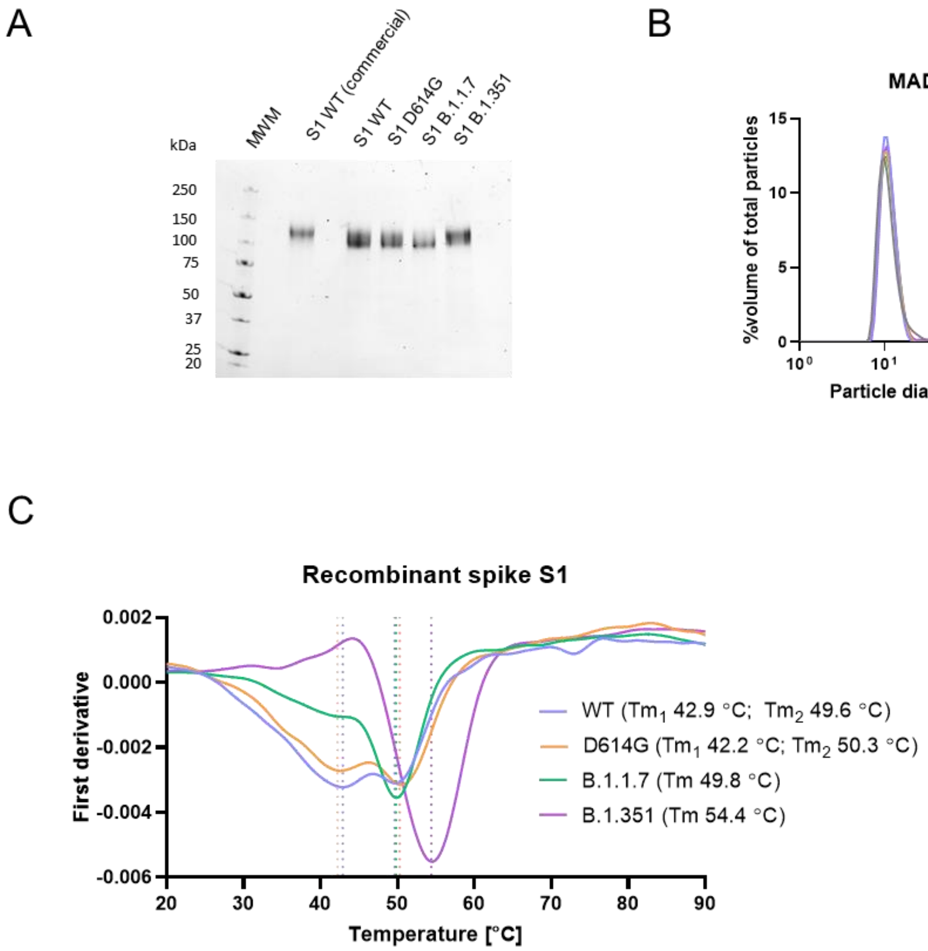


**Supplementary Figure 1. Characterisation of SARS-CoV-2 S1 variants**

A) SDS-PAGE of SARS-CoV-2 S1 purified proteins from WT, D614G, B.1.1.7 and B.1.351 variants, in comparison to a commercially sourced recombinant S1 protein. All proteins showed >95% purity. B) Particle size distribution of recombinant purified SARS.-CoV-2 S1 variants analysed via MADLS. C) Thermal stability analysis of purified SARS-CoV-2 variants showing increased Tm for B.1.1.7 and B.1.351 variants compared to WT S1.

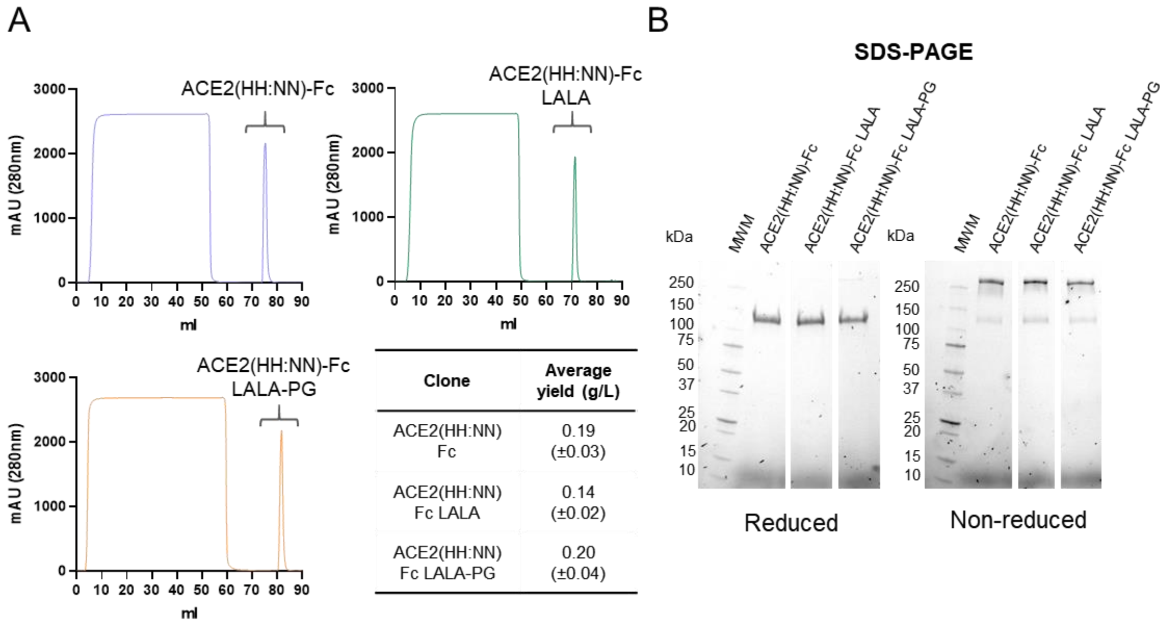


| Clone | Average yield (g/L) |
|---|---|
| ACE2(HH:NN) Fc | 0.19 (±0.03) |
| ACE2(HH:NN) Fc LALA | 0.14 (±0.02) |
| ACE2(HH:NN) Fc LALA-PG | 0.20 (±0.04) |

**Supplementary figure 2. Purification profile of ACE2(HH:NN)-Fc variants**

A) Representative protein A purification profile of ExpiCHO expressed ACE2(HH:NN)-Fc (blue), ACE2(HH:NN)-Fc LALA (green) and ACE2(HH:NN)-Fc LALA-PG (orange) fusion proteins using AKTA pure and Mab select PrismA columns with in-line HiTrap desalting column. n=4, n=3, n=10, respectively; Mean (±SD). B) Representative SDS-PAGE of purified ACE2(HH:NN)-Fc, ACE2(HH:NN)-Fc LALA and ACE2(HH:NN)-Fc LALA-PG in reducing and non-reducing conditions showing > 95% purity.

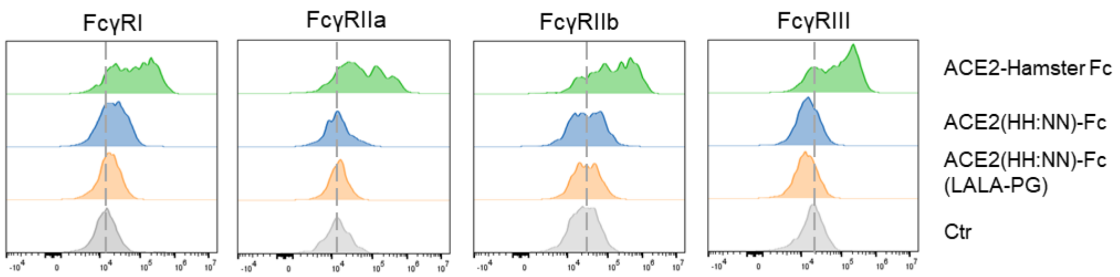


**Supplementary figure 3. Hamster FcyR binding**

Fc-mediated binding capacity of ACE2(HH:NN) WT Fc (blue), LALA-PG Fc (orange) or ACE2(HH:NN) hamster Fc (green) to HEK293T cells expressing hamster FcγRI, FcγRIIa, FcγRIIb and FcγRIII receptors, detected with biotinylated SARS-CoV-2 S1 and streptavidin conjugated secondary agent. No binding was detected with ACE2(HH:NN)-Fc constructs carrying the LALA-PG mutations on the hamster FcγRs, while limited binding was detected with the ACE2(HH:NN) WT Fc.

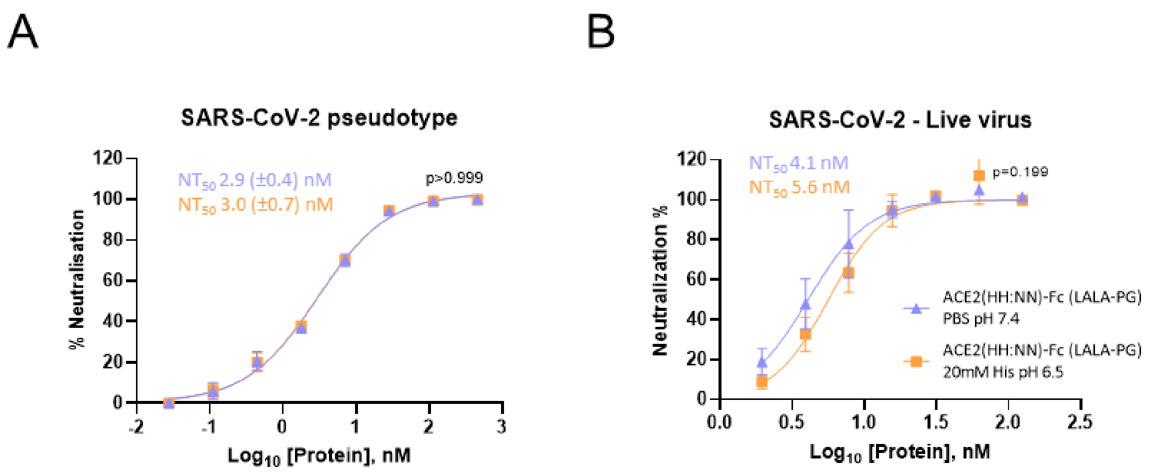


**Supplementary figure 4. Comparison of neutralisation efficiency**

Neutralisation assay of SARS-CoV-2 pseudotype (A) or live virus (B) with ACE2(HH:NN)-Fc LALA-PG in PBS pH 7.4 (blue) and 20 mM His pH 6.5 (orange). Both formulations show comparable neutralisation efficiencies (Mean ± SD). Unpaired t test of AUC (A t=0.000, df=32; B t=1.317, df=28).

**Supplementary Table 1 – Cell microarray antigens**

Xlsx table in attachment file
